# Pathological claustrum activity drives aberrant cognitive network processing in human chronic pain

**DOI:** 10.1101/2023.11.01.564054

**Authors:** Brent W. Stewart, Michael L. Keaser, Hwiyoung Lee, Sarah M. Margerison, Matthew A. Cormie, Massieh Moayedi, Martin A. Lindquist, Shuo Chen, Brian N. Mathur, David A. Seminowicz

**Author notes:** <u>Declaration of Interests</u>: The authors declare no competing interests. Correspondence should be addressed to: Brian N. Mathur HSF III 9179 670 West Baltimore Street Baltimore, MD 21201 and David A. Seminowicz Robarts Research Institute 100 Perth Dr London, ON N6A 5K8.

## Abstract

Aberrant cognitive network activity and cognitive deficits are established features of chronic pain. However, the nature of cognitive network alterations associated with chronic pain and their underlying mechanisms require elucidation. Here, we report that the claustrum, a subcortical nucleus implicated in cognitive network modulation, is activated by acute painful stimulation and pain-predictive cues in healthy participants. Moreover, we discover pathological activity of the claustrum and a lateral aspect of the right dorsolateral prefrontal cortex (latDLPFC) in migraine patients. Dynamic causal modeling suggests a directional influence of the claustrum on activity in this latDLPFC region, and diffusion weighted imaging (DWI) verifies their structural connectivity. These findings advance understanding of claustrum function during acute pain and provide evidence of a possible circuit mechanism driving cognitive impairments in chronic pain.

## Introduction

Pain demands attention (Eccleston & Crombez, 1999). Simultaneous acute pain and cognitive processing appear to compete for resources in the brain, with increased pain associated with reduced cognitive task performance and increased cognitive load associated with diminished pain (Buhle & Wager, 2010). Patients with chronic pain often report cognitive difficulties, and significant impairments in executive function are detectable in patients from a variety of chronic pain conditions (Baker et al., 2016; Berryman et al, 2014; Landrø et al., 2013). Cognitive impairments are linked with cortical network dysfunction in multiple neuropsychiatric disorders (Bassett et al., 2008; Rombouts et al., 2005; Yao et al., 2010), including chronic pain, which is known to alter cortical network activity (Baliki et al., 2008; Hashmi et al., 2013). Indeed, experimental pain, in an intensity-dependent fashion, increases co-activation of brain regions that respond to a cognitive task in a difficulty-dependent fashion (Seminowicz & Davis, 2007), and when patients with chronic pain perform a cognitive task, they exhibit greater activity across the brain relative to controls, comprising reduced areas of deactivation and wider areas of activation (Ceko et al., 2015; Mathur et al., 2015; Seminowicz et al., 2011). How cortical network disruptions and associated cognitive impairments arise in chronic pain is unclear.

The claustrum is a subcortical nucleus implicated in orchestrating cortical networks. Studies of rodent claustrum circuitry demonstrate that the claustrum preferentially routes input from frontal cortices to numerous output cortical regions involved in higher order processing (Qadir et al., 2018; Qadir et al., 2022; White et al., 2017; White & Mathur, 2018a; White & Mathur, 2018b). Human imaging reveals that the claustrum shares functional connectivity with multiple canonical resting state networks (Krimmel et al., 2019), possesses the greatest anatomical connectivity in the brain by volume, and is a critical hub in global cortical network architecture (Torgerson et al., 2015). Functionally, in animal models, claustrum activation is required for optimal performance in cognitively demanding tasks (Atlan et al., 2018; White et al., 2018; White et al., 2020), and in humans, significant claustrum activation occurs at the onset of a difficult cognitive task, when cognitive network activity appears (Krimmel et al., 2019). These findings led us to propose that the claustrum instantiates cortical networks for cognitive control (Madden et al., 2022). Because there is evidence that the claustrum responds to acute pain (Paul et al., 2021), and pain is a cognitive load, it is possible that pain induces cognitive control network formation through claustrum activation. Additionally, this process may be altered in chronic pain, where networks and cognition are dysfunctional.

Testing these ideas, we measured claustrum signal using fMRI during experimental pain, pain anticipation, cognitive conflict, and resting-state conditions in samples of healthy controls and patients with chronic pain (migraine). Our findings indicate that the claustrum is responsive to pain onset or pain-predictive cues, that migraine patients recruit altered cognitive task networks characterized by the addition of a pain-sensitive brain region, and that migraine patients exhibit pathological claustrum activity that drives chronic pain-associated cognitive network alterations. These findings support the notion that claustrum dysfunction may underlie aberrant cognitive network processing observed in chronic pain.

## Results

### Claustrum BOLD signal increases in response to experimental heat pain

We first predicted that the claustrum activates at the onset of acute pain in healthy conditions, analogously to its activation at the onset of a difficult cognitive task (Krimmel et al., 2019). Dataset 1 included Blood Oxygenation Level Dependent (BOLD) fMRI (TR = 2.5 seconds) pain scans from n = 34 healthy participants. Data were acquired from one (n = 10), two (n = 6), or three (n = 18) imaging sessions depending on participant availability, and two pain runs (8 minutes/run) were acquired from each participant per imaging session. Each run consisted of five trials of thermal stimulation applied to the left forearm. All trials began with a two second temperature ramp up from 32°C (baseline) to a subject-specific non-painful warm temperature (8°C less than the subject’s painful temperature), which was held for 28 seconds. Another two second ramp up directly followed the warm stimulation, this time arriving at a temperature rated by the subject as moderately painful (5-7 on a 0-10 numeric rating scale). The painful stimulus was also held for 28 seconds before the trial ended with a two second ramp down to baseline temperature (Fig. 1B).

**Figure 1.**
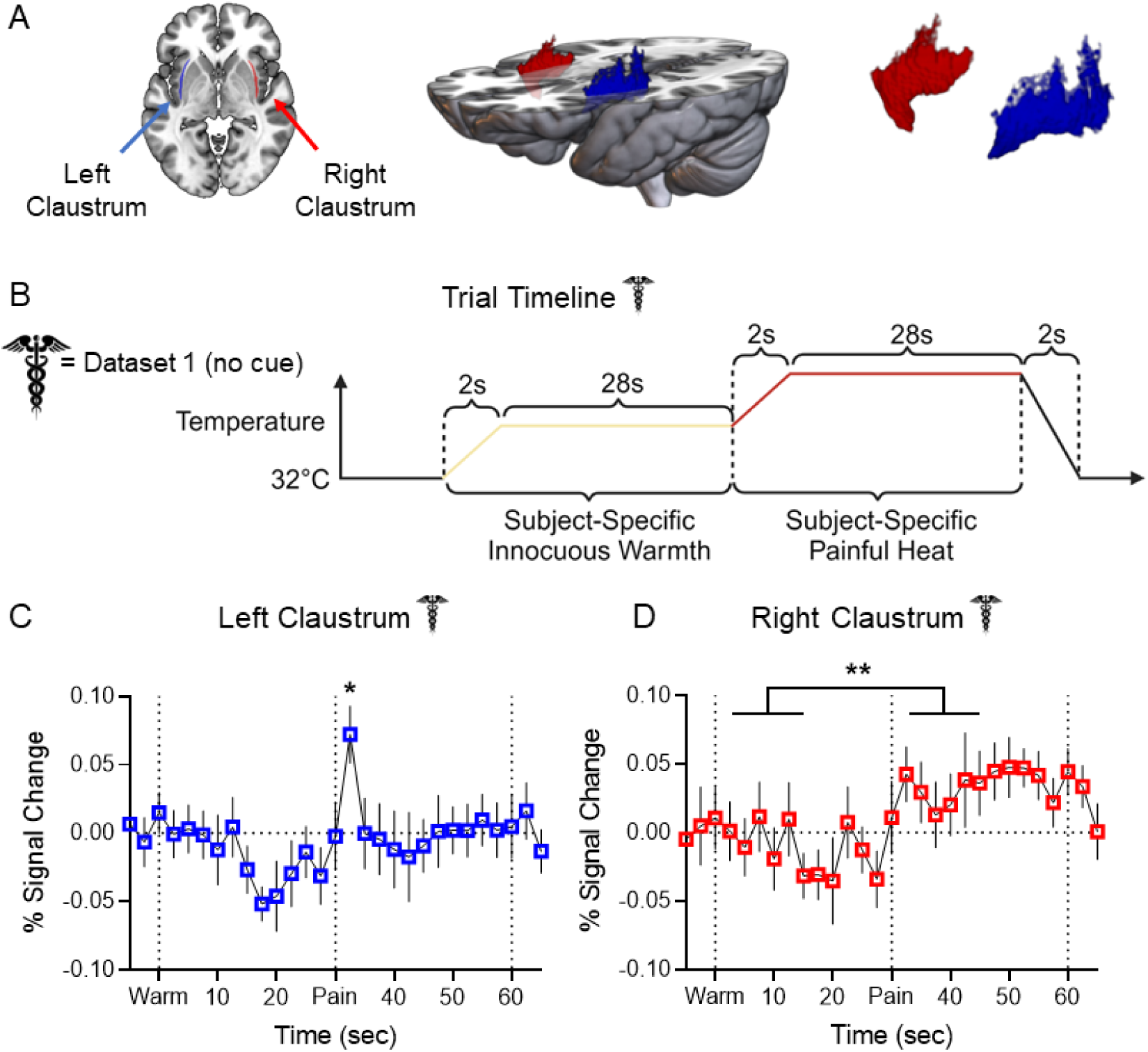
Claustrum BOLD signal increases in response to experimental heat pain. **(A) Left:** Horizontal slice of MNI template displaying LCL (blue) and RCL (red) in neurological orientation. **Middle:** 3D rendering of MNI template displaying dorsoventral extent and placement of LCL and RCL. **Right:** 3D rendering displaying shape of LCL and RCL. **(B)** Experimental timeline for each trial in Dataset 1 pain scans. **(C)** Average FIR timeseries of LCL BOLD percent signal change showed a significant increase at t = 32.5 sec (pain “onset”: t = 3.011, p-FDR = 0.023) in Dataset 1 healthy participants (n = 34). **(D)** Average FIR timeseries of RCL BOLD percent signal change showed significantly greater signal during pain than warm timepoints (t = 2.882, p-FDR = 0.004). Error bars show standard error of the mean across subjects. * represents a p-value of < 0.05, ** < 0.01, *** < 0.001. Caduceus symbol signifies Dataset 1 results. Image by Clker-Free-Vector-Images from Pixabay. Unfilled bars/plot points signify healthy participant results. Acronyms: ALFF – Amplitude of Low Frequency Fluctuations, BOLD – Blood Oxygenation Level Dependent, DCM – Dynamic Causal Modeling, DWI – Diffusion Weighted Imaging, FIR – Finite Impulse Response, GLM – General Linear Model, HCP – Human Connectome Project, HRF – Hemodynamic Response Function, LaINS – Left Anterior Insula, latDLPFC – Lateral Right Dorsolateral Prefrontal Cortex (see text), LCL – Left Claustrum, LInsFl – Left Insular Flank (see text), LV – Latent Variable, medDLPFC – Medial Right Dorsolateral Prefrontal Cortex (see text), MNI – Montreal Neurological Institute, MSIT – Multi-Source Interference Task, RaINS – Right Anterior Insula, RCL – Right Claustrum, RInsFl – Right Insular Flank (see text), TR – Repetition Time, WM – White Matter. Figures generated in GraphPad Prism, MRICroGL, and BioRender.com.

A General Linear Model (GLM) using a Finite Impulse Response (FIR) to model the hemodynamic response function (HRF) was fitted to generate timeseries of claustrum BOLD percent signal change. Timeseries were averaged across trials within subjects and then across the group. Figure 1 illustrates the average BOLD percent signal change in the left (LCL; Fig. 1C) and right (RCL; Fig. 1D) claustrum with error bars representing standard error of the mean across subjects. The start of the warm ramp up (t = 0 seconds) was defined as w0, and the start of the pain ramp up (t = 30 seconds) was defined as p0. To limit comparisons and prevent selection bias, statistical models included four timepoints from each stimulus. Accounting for hemodynamic delay, “onset” timepoints were defined as w1-3 and p1-3, and “block” timepoints were defined as w6 and p6, the midpoint of each stimulation block. Linear mixed effects models compared the percent signal change at each of these timepoints to the average pre-trial (t-2 and t-1) percent signal change.

As predicted, LCL signal exhibited a significant increase over pre-stimulus baseline at pain onset (p1: t = 3.011, false discovery rate corrected p-FDR = 0.023). RCL did not exhibit significant signal change at any timepoint compared to baseline. However, a linear mixed effects model assessing condition effects demonstrated a significant effect of pain greater than warm (t = 2.882, p-FDR = 0.004), such that RCL signal during painful stimulation timepoints was generally greater than warm stimulation timepoints. LCL and RCL signal changes were not attributable to subject motion (Supplemental Fig. S1A).

### Claustrum BOLD signal is distinguishable from neighboring regions

Because aspects of the claustrum are thinner than the 2mm isomorphic functional image voxels analyzed in this study, claustrum region of interest (ROI) analyses are vulnerable to partial volume effects. We therefore implemented Small Region Confound Correction (SRCC; Barrett et al., 2020; Krimmel et al., 2019) for all claustrum analyses (see Methods). To test if obtained claustrum signal was distinguishable from proximal structures, we measured responses in ROIs comprising a medial portion of each hemisphere’s insular cortex that “flank” the claustrum (Fig. 2A: left insular flank [LInsFl] – light blue, right insula flank [RInsFl] – pink; see Methods). FIR timeseries of bilateral flanking insular cortex ROIs illustrate BOLD signal increases of greater amplitude and longer duration than in LCL or RCL at pain onset (t = 30) and pain offset (t = 60) (Fig. 2B-C).

**Figure 2.**
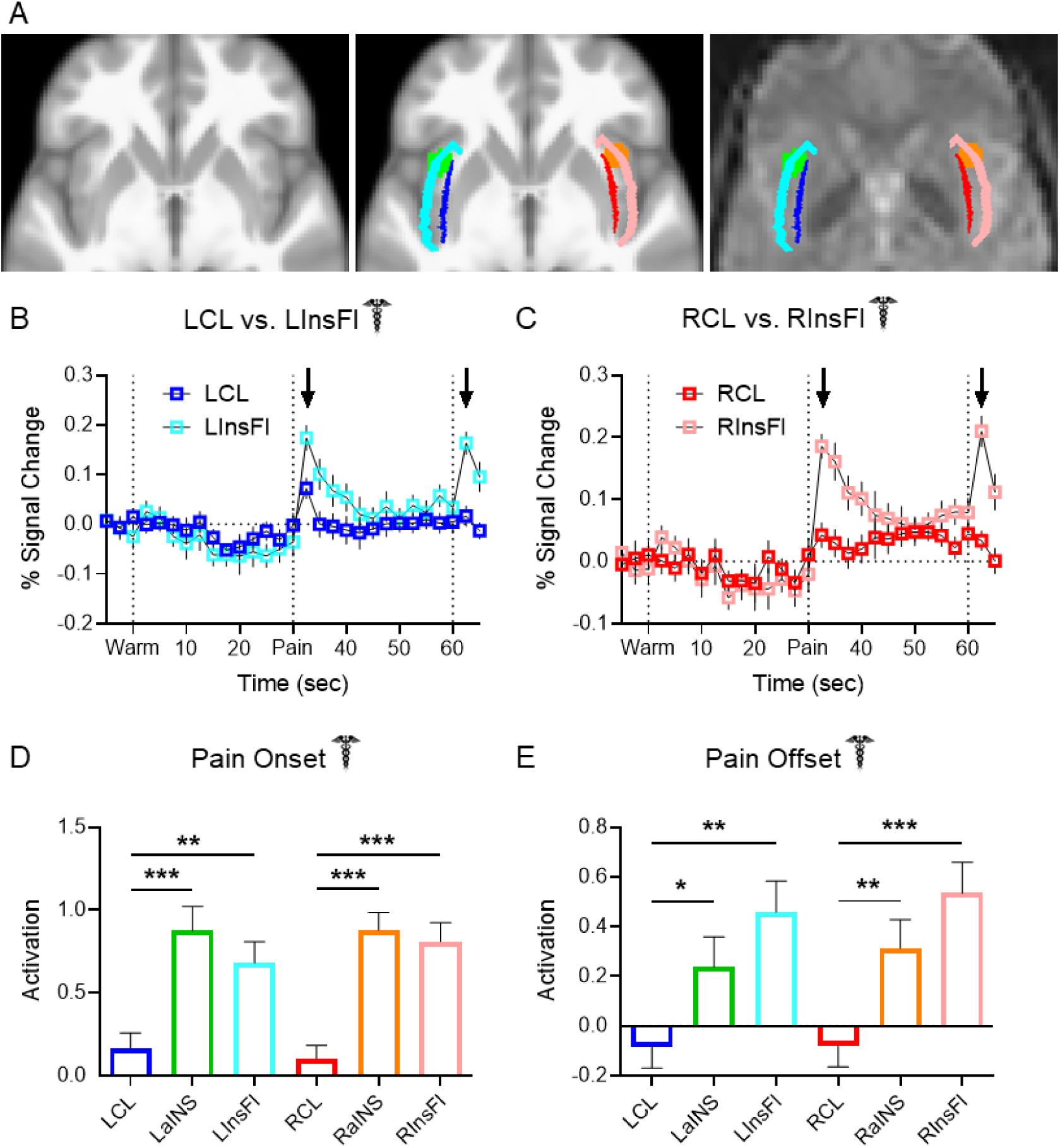
Claustrum BOLD signal is distinguishable from neighboring regions. **(A) Left:** Horizontal slice of average anatomical scan from 56 Dataset 1 subjects. **Middle:** Same slice as left depicting shapes and locations of LCL (blue), LInsFl (light blue), LaINS (green), RCL (red), RInsFl (pink), and RaINS (orange) ROIs. **Right:** Same z-stack as left and middle with ROIs overlaid on a Dataset 1 healthy participant’s preprocessed functional resting state image. **(B – C)** Average FIR timeseries of Dataset 1 healthy participant thermal stimulation trials displayed greater increases at pain onset and pain offset (black arrows) in (B) LInsFl vs. LCL and (C) RInsFl vs. RCL BOLD percent signal change. No statistical tests were performed on (B) & (C). **(D)** Regional activation at pain onset was compared to assess the distinctiveness of claustrum signal change to a stimulus predicted to evoke similar responses across proximal ROIs. Average parameter estimation (regression slope) detected significantly different activation at pain onset (i.e., first two seconds of thermal stimulation) between LCL and LaINS (t = 4.086, p-FDR < 0.001), LCL and LInsFl (t = 3.266, p-FDR = 0.002), RCL and RaINS (t = 5.816, p-FDR < 0.001), and RCL and RInsFl (RCL vs. RInsFl: t = 5.039, p-FDR < 0.001). **(E)** Regional activation at pain offset was compared upon observing divergent claustrum and insular flank timeseries at pain offset in (B) & (C) to assess potential functional differences between proximal ROIs. Significantly different activation at pain offset was observed between LCL and LaINS (t = 2.133, p-FDR = 0.037), LCL and LInsFl (t = 3.551, p-FDR = 0.001), RCL and RaINS (t = 2.773, p-FDR = 0.0096), and RCL and RInsFl (t = 4.095, p-FDR < 0.001). Assessment of region effects via unpaired t-tests was limited to comparisons of LCL vs. LaINS, LCL vs. LInsFl, RCL vs. RaINS, and RCL vs. RInsFl.

To facilitate direct comparisons between regions, singular activation values were derived from GLM analyses in SPM12 (http://www.fil.ion.ucl.ac.uk/spm/software/spm12/). Unlike the previous models using FIR, these analyses assumed a canonical HRF and required further dividing experimental trials into component conditions. Warm and pain stimulation conditions were separated into two second “onset” conditions corresponding to the temperature ramp up and 28 second “block” conditions corresponding to the remainder of the stimulation. Bilateral anterior insular cortices (LaINS & RaINS; Fig. 2A green and orange, respectively) are major Salience Network (Seeley et al., 2007; Taylor et al., 2009) regions and were also predicted to respond to pain. LaINS and RaINS ROIs were therefore analyzed as positive controls and to further test the distinctiveness of claustrum BOLD signal.

Unpaired t-tests revealed claustrum signal at pain onset to be significantly different than anterior insula and flanking insula signal in both hemispheres (Fig. 2D, LCL vs. LaINS: t = 4.086, p-FDR < 0.001; LCL vs. LInsFl: t = 3.266, p-FDR = 0.002; RCL vs. RaINS: t = 5.816, p-FDR < 0.001; RCL vs. RInsFl: t = 5.039, p-FDR < 0.001).

Claustrum responses were also distinguishable from neighboring regions at pain offset (Fig. 2E). Unpaired t-tests detected significantly different activation between LCL and LaINS (t = 2.133, p-FDR = 0.037), LCL and LInsFl (t = 3.551, p-FDR = 0.001), RCL and RaINS (t = 2.773, p-FDR = 0.0096), and RCL and RInsFl (t = 4.095, p-FDR < 0.001).

### Claustrum BOLD signal increases in response to a pain-predictive cue

An additional dataset (Dataset 2) allowed investigation of claustrum effects in more conditions. In this experiment, healthy participants (n = 39) underwent a single BOLD fMRI imaging session comprising as many as five runs (8 minutes/run; TR = 1.75 secs) in which they experienced pseudorandom durations and intensities of thermal stimulation to their lower left leg (10 trials/run). Intensities included non-painful warmth (38°C), “moderate” pain defined as in Dataset 1, “slight” pain, and “intense” pain, defined as 1°C less than or greater than “moderate” respectively. Notably, each trial began with an auditory “doorbell sound” cue and a 7.5 second anticipation period prior to thermal stimulation (Fig. 3A). This dataset consequently allowed the analysis of claustrum responses to a pain-predictive stimulus.

**Figure 3.**
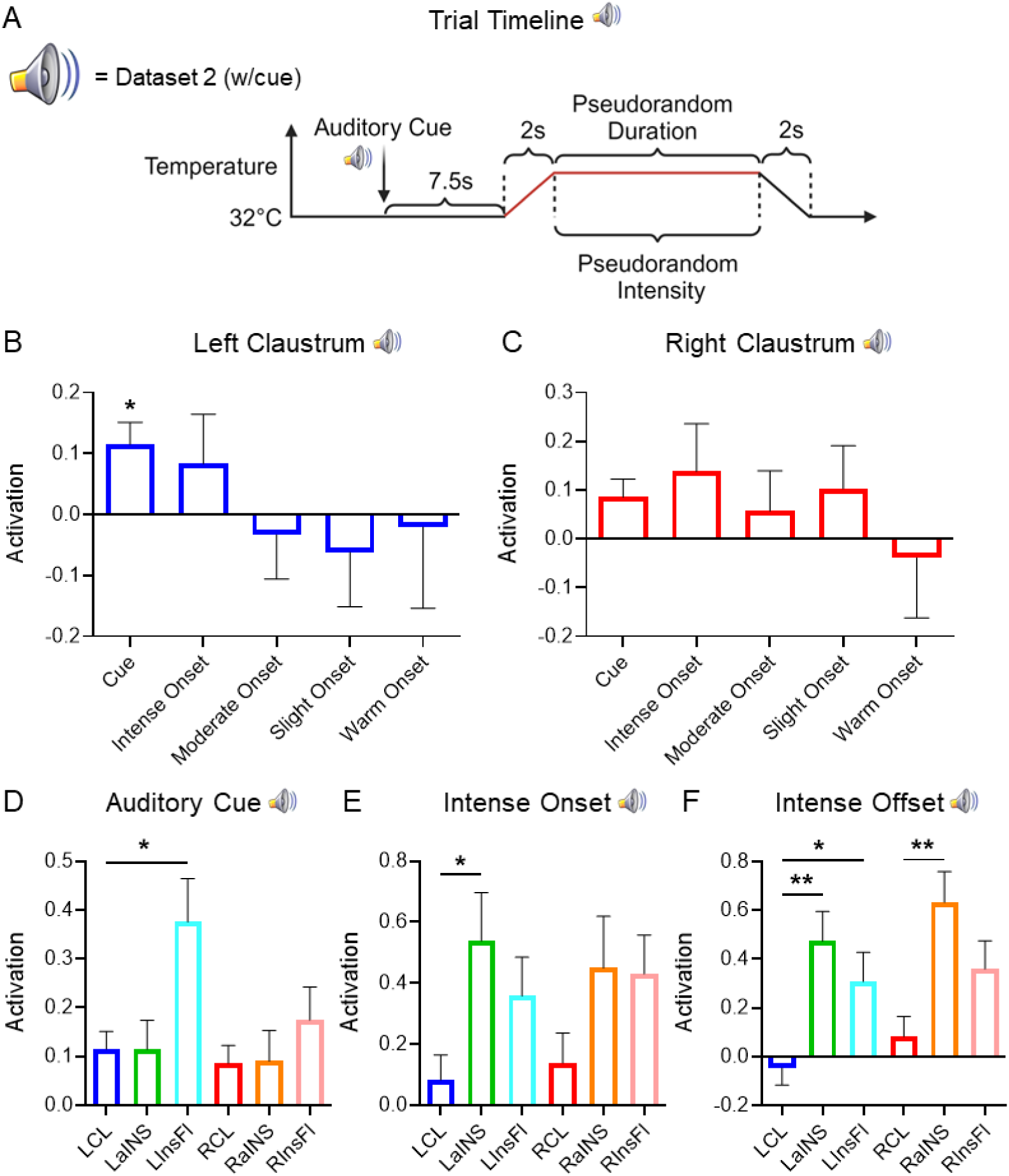
Claustrum BOLD signal increases in response to a pain-predictive cue. **(A)** Timeline of thermal stimulation for each trial in Dataset 2, which comprised (n = 39) healthy participants. **(B)** LCL displayed significant activation in response to the cue preceding thermal stimulation (t = 3.172, p-FDR = 0.030) but not to the onset of any thermal stimulation conditions following the cue (intense onset: t = 1.034, p-FDR = 0.615; moderate onset: t = 0.460, p-FDR = 0.811; slight onset: t = 0.699, p-FDR = 0.699; warm onset: t = 0.153, p-FDR = 0.879). **(C)** RCL did not display significant activation in response to the cue (t = 2.341, p-FDR = 0.123) or the onset of any thermal stimulation conditions (intense onset: t = 1.416, p-FDR = 0.550; moderate onset: t = 0.707, p-FDR = 0.699; slight onset: t = 1.176, p-FDR = 0.615; warm onset: t = 0.319, p-FDR = 0.835). **(D)** Significantly different activation in response to the auditory cue was observed between LCL and LInsFl (t = 2.713, p-FDR = 0.033). Significantly different activation in response to the auditory cue was not observed between LCL and LaINS (t = 0.009, p-FDR = 0.993), RCL and RaINS (t = 0.059, p-FDR = 0.993), or RCL and RInsFl (t = 1.161, p-FDR = 0.499). **(E)** In response to the onset of intense pain, significantly different activation was observed between LCL and LaINS (t = 2.564, p-FDR = 0.049), but not between LCL and LInsFl (t = 1.817, p-FDR = 0.098), RCL and RaINS (t = 1.587, p-FDR = 0.117), or RCL and RInsFl (t = 1.816, p-FDR = 0.098). **(F)** In response to the offset of intense pain, significantly different activation was observed between LCL and LaINS (t = 3.745, p-FDR = 0.001), LCL and LInsFl (t = 2.553, p-FDR = 0.017), RCL and RaINS (t = 3.637, p-FDR = 0.001), but not between RCL and RInsFl (t = 1.938, p-FDR = 0.056). Assessment of condition effects in (B) & (C) via unpaired t-tests was limited to comparisons of cue with thermal stimulation onset conditions. Assessment of region effects via unpaired t-tests in (D) – (F) was limited to comparisons of LCL vs. LaINS, LCL vs. LInsFl, RCL vs. RaINS, and RCL vs. RInsFl. Speaker symbol signifies Dataset 2 results. Image by Clker-Free-Vector-Images from Pixabay.

One sample t-tests of Dataset 2 results revealed significant activation in LCL (Fig. 3B, Cue: t = 3.172, p-FDR = 0.030) but not RCL (Fig. 3C, Cue: t = 2.341, p-FDR = 0.123) in response to the pain-predictive auditory cue. Bilateral claustrum regions did not exhibit significant activation in response to the onset of any thermal stimulation condition when preceded by an auditory cue (Fig. 3B-C, LCL intense onset: t = 1.034, p-FDR = 0.615; LCL moderate onset: t = 0.460, p-FDR = 0.811; LCL slight onset: t = 0.699, p-FDR = 0.699; LCL warm onset: t = 0.153, p-FDR = 0.879; RCL intense onset: t = 1.416, p-FDR = 0.550; RCL moderate onset: t = 0.707, p-FDR = 0.699; RCL slight onset: t = 1.176, p-FDR = 0.615; RCL warm onset: t = 0.319, p-FDR = 0.835).

LCL signal was significantly different than LInsFl in response to the cue (Fig. 3D, t = 2.713, p-FDR = 0.033). Significant differences were not identified when comparing cue responses between LCL and LaINS (t = 0.009, p-FDR = 0.993), RCL and RaINS (t = 0.059, p-FDR = 0.993), or RCL and RInsFl (t = 1.161, p-FDR = 0.499).

In response to the onset of intense pain, LCL signal was significantly different than LaINS (Fig. 3E, t = 2.564, p-FDR = 0.049). Significant differences were not detected when comparing LCL and LInsFl (t = 1.817, p-FDR = 0.098), RCL and RaINS (t = 1.587, p-FDR = 0.117), or RCL and RInsFl (t = 1.816, p-FDR = 0.098).

Like Dataset 1, claustrum, insula flank, and anterior insula ROI signals diverged at the offset of intense pain in Dataset 2 (Fig. 3F). LCL signal at intense offset was significantly different from LaINS (t = 3.745, p-FDR = 0.001) and LInsFl (t = 2.553, p-FDR = 0.017). RCL signal was significantly different from RaINS (t = 3.637, p-FDR = 0.001) while exhibiting a trending difference with RInsFl (t = 1.938, p-FDR = 0.056).

### Migraine patients exhibit greater cognitive task-associated network activity than controls

Upon establishing claustrum responses to painful stimulation or pain cues in healthy participants, we extended our investigation to a sample of patients with chronic pain. Due to previous reports of altered cognitive task-associated activity in multiple chronic pain conditions (Ceko et al., 2015; Mathur et al., 2015; Seminowicz et al., 2011) we tested if an altered cognitive task network phenotype was present in Dataset 1, which included BOLD fMRI cognitive task scans from healthy controls (n = 35) and migraine patients (n = 112).

During cognitive task scans, participants performed the multi-source interference task (MSIT; Bush et al., 2003). The task consisted of 20 second blocks of consecutive trials representing one of three conditions. One condition was an easy cognitive task. In each trial, participants were presented with an array of three numbers in which one number was different than the other two. The participant was directed to press one of three buttons on a response pad corresponding to the unique number, and in the easy condition the unique number matched its position in the array (e.g., “1-3-3”, “1-2-1”, “2-2-3”). The second condition was a difficult cognitive task which was structured similarly to the easy condition except the unique number did not match its position in the array (e.g., “3-1-1”, “2-1-2”, “3-3-2”). The third condition was a motor control in which an asterisk appeared on the screen in one of three positions (e.g., “* – –”, “– * –”, “– – *”) and the participant pressed a button corresponding to the position of the asterisk.

Partial Least Squares (PLS; McIntosh et al., 1996) yields “latent variables” (LVs) representing multivariate patterns of brain activity that covary with experimental conditions. PLS analyses of Dataset 1 whole-brain cognitive task-associated activity identified significant LVs sensitive to task difficulty comprised of signal increases resembling Fronto-Parietal Network and Cingulo-Opercular Network motifs (Dosenbach et al., 2007; Dosenbach et al., 2008) as well as signal decreases resembling the Default Mode Network (DMN; Greicius et al., 2003) within healthy controls (Fig. 4A - top left, n = 35, LV1: p < 0.001) and migraine patients (Fig. 4A - top right, n = 112, LV1: p < 0.001). The overlap of regions demonstrating cognitive task-associated signal increases in these LVs indicated a wider spatial extent of activation in patients than controls (Fig. 4A - bottom left). A group effects PLS analysis identified a pattern of brain regions where cognitive task-associated activity significantly differed between groups (Fig. 4A - bottom right, LV1: p < 0.048). A cluster report of this LV identified 12 significantly different clusters, all of which exhibited greater activity in patients than controls (Supplemental Table 1).

**Figure 4.**
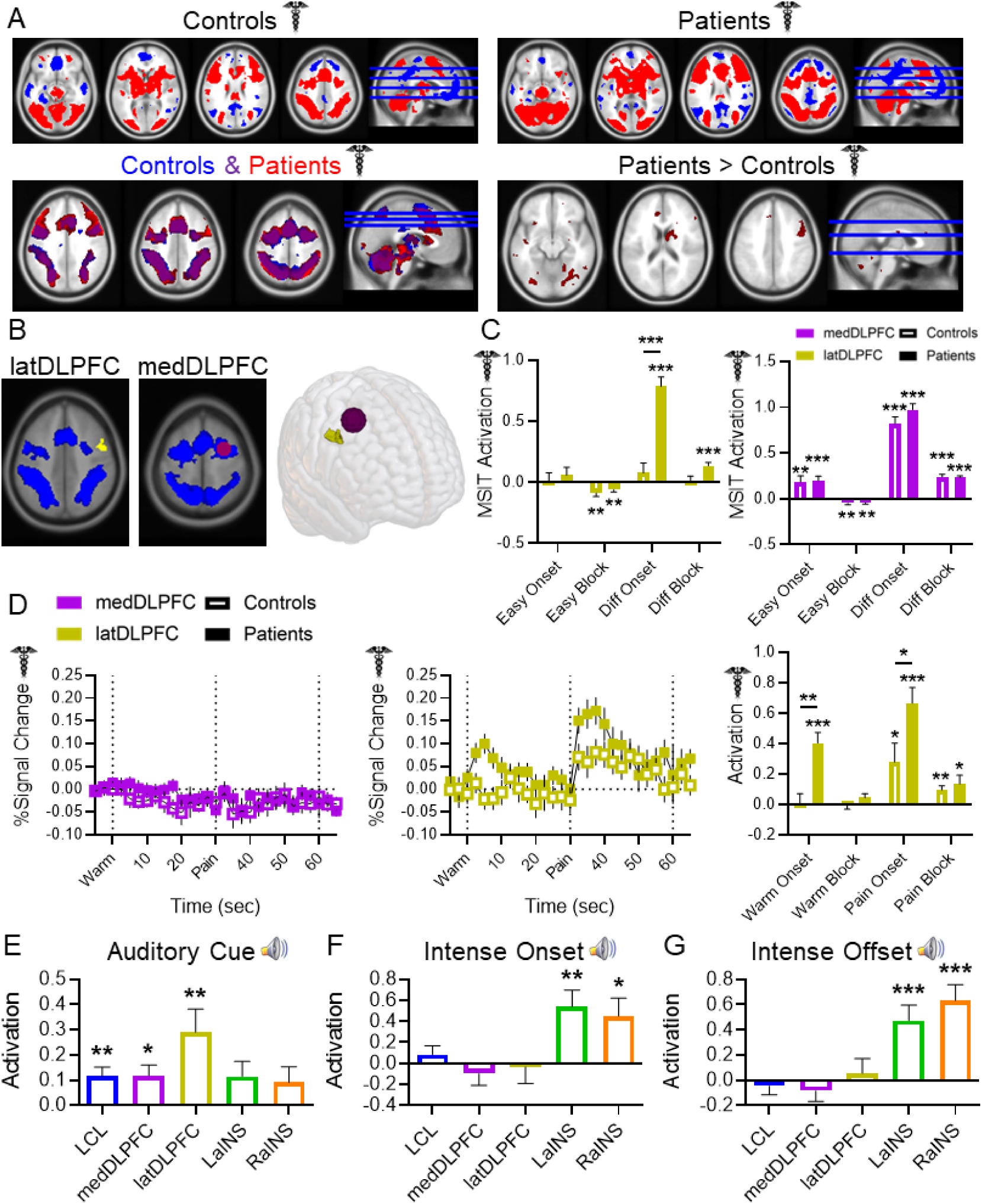
Migraine patients engage a pain-responsive prefrontal cortex region during pain-free cognitive task processing. **(A) Top left:** Horizontal slices with reference sagittal slice illustrating the multivariate pattern accounting for the most variance (LV1) in brain activity associated with cognitive task conditions in Dataset 1 healthy controls (n = 35, LV1: p < 0.001). **Top right:** LV1 of brain activity associated with cognitive task conditions in Dataset 1 migraine patients (n = 112, LV1: p < 0.001). All voxels shown in top row exhibited signal increases (red) or decreases (blue) with increasing cognitive load (motor control tapping condition, easy cognitive task, difficult cognitive task). **Bottom left:** Voxels exhibiting significant signal increases in LV1 of healthy controls (blue), LV1 of migraine patients (red), and their overlap (purple). **Bottom right:** Clusters (> 50 voxels) depicting significantly different cognitive task-associated signal changes between patients and controls (patients vs. controls LV1: p < 0.048). All significant clusters exhibited greater signal in patients than controls. **(B) Left:** latDLPFC (gold) depicted alongside healthy controls’ cognitive task LV1 (blue). Note how latDLPFC fell entirely outside the healthy control cognitive task network. **Middle:** medDLPFC (purple) overlaid on healthy controls’ cognitive task LV1 (blue). medDLPFC (center: 28, 0, 52; radius: 10mm) was centered within the healthy control cognitive task LV1 and the migraine patient cognitive task LV1 (not pictured). **Right:** 3D rendering depicting shapes and locations of latDLPFC and medDLPFC. **(C) Left:** Cognitive task-induced latDLPFC activation was only observed in patients during difficult onset (t = 10.129, p-FDR < 0.001) and difficult block (t = 5.178, p-FDR < 0.001) conditions. Two-way ANOVA found significant main effects of group (F (1, 580) = 20.38, p < 0.001), condition (F (3, 580) = 18.84, p < 0.001), and their interaction (F (3, 580) = 10.20, p < 0.001), and post-hoc comparisons detected significantly greater activation in patients than controls during the difficult onset condition (t = 7.003, p < 0.001). **Right:** Significant medDLPFC activation was detected in cognitive task conditions in healthy controls (easy onset: t = 2.835, p-FDR = 0.008; difficult onset: t = 11.328, p-FDR < 0.001; difficult block: t = 9.327, p-FDR < 0.001) and migraine patients (easy onset: t = 3.956, p-FDR < 0.001; difficult onset: t = 14.767, p-FDR < 0.001; difficult block: t = 11.980, p-FDR < 0.001). No main effect of group was detected (F (1, 580) = 0.939, p = 0.333). **(D) Left:** Average FIR timeseries of medDLPFC BOLD percent signal change during thermal stimulation in healthy controls (n = 34) and migraine patients (n = 105). **Middle:** Average FIR timeseries of latDLPFC BOLD percent signal change during thermal stimulation in both groups. No statistical tests were performed on these timeseries. **Right:** Significant latDLPFC activation was observed in both groups at pain onset (controls: t = 2.265, p-FDR = 0.048; patients: t = 6.481, p-FDR < 0.001) and during pain block (controls: t = 3.557, p-FDR = 0.003; patients: t = 2.358, p-FDR = 0.041). Two-way ANOVA found significant main effects of group (F (1, 548) = 11.38, p < 0.001), and condition (F (3, 548) = 8.985, p < 0.001), but not their interaction (F (3, 548) = 2.396, p = 0.067), and post hoc comparisons detected significantly greater activation in patients than controls during the warm onset (t = 3.08, p = 0.009) and pain onset (t = 2.973, p = 0.012) conditions. **(E)** medDLPFC and latDLPFC activation aligned with LCL but diverged from bilateral anterior insula in Dataset 2 in response to the auditory cue (LCL: t = 3.172, p-FDR = 0.008; medDLPFC: t = 2.811, p-FDR = 0.013; latDLPFC: t = 3.151, p-FDR = 0.008; LaINS: t = 1.929, p-FDR = 0.077; RaINS: t = 1.437, p-FDR = 0.159), **(F)** the onset of intense pain (LCL: t = 1.034, p-FDR = 0.470; medDLPFC: t = 0.896, p-FDR = 0.470; latDLPFC: t = 0.216, p-FDR = 0.830; LaINS: t = 3.409, p-FDR = 0.008; RaINS: t = 2.649, p-FDR = 0.029), and **(G)** the offset of intense pain (LCL: t = 0.678, p-FDR = 0.627; medDLPFC: t = 1.025, p-FDR = 0.519; latDLPFC: t = 0.483, p-FDR = 0.632; LaINS: t = 3.926, p-FDR < 0.001; RaINS: t = 5.003, p-FDR < 0.001). Unfilled bars/plot points signify healthy participant results. Filled bars/plot points signify migraine patient results.

### Migraine patients engage a pain-responsive prefrontal cortex region during pain-free cognitive task processing

In light of our hypothesis that claustrum initiates cognitive networks, the observation of altered cognitive network recruitment in patients motivated deeper investigation of differentially activated cognitive network nodes.

The peak voxel exhibiting significantly greater cognitive task-associated activity in patients than controls (Fig. 4A - bottom right; Supplemental Table 1, cluster 1, peak voxel: (52, 0, 44)) fell within a region highlighted in red in the bottom left panel of Figure 4A, meaning it displayed cognitive task-associated signal increases only in patients. We therefore asked what conditions this voxel and its cluster respond to in healthy controls, and what additional function this region may perform in patients during a cognitive task. A contiguous portion of the cluster containing only voxels outside the healthy control cognitive task-associated regions (96 voxels) was selected for further analysis. The ROI was within the right dorsolateral prefrontal cortex (DLPFC), but it is designated herein as latDLPFC (Fig. 4B - left). Another region, falling within both groups’ respective cognitive task-associated networks and exhibiting no significant cognitive task activity differences between groups was selected for comparison. This ROI (center: (28, 0, 52); radius: 10mm) was also within the right DLPFC but was more dorsal and medial than latDLPFC. It is designated herein as medDLPFC (Fig. 4B - middle).

For consistency with pain analyses, each MSIT condition was separated into an “onset” portion consisting of the condition’s first two seconds and a “block” portion consisting of the condition’s remaining 18 seconds. As expected based on PLS findings, according to one sample t-tests (Fig. 4C - right), medDLPFC exhibited significant activation in response to cognitive task conditions in both groups (controls easy onset: t = 2.835, p-FDR = 0.008; controls difficult onset: t = 11.328, p-FDR < 0.001; controls difficult block: t = 9.327, p-FDR < 0.001; patients easy onset: t = 3.956, p-FDR < 0.001; patients difficult onset: t = 14.767, p-FDR < 0.001; patients difficult block: t = 11.980, p-FDR < 0.001), and a two-way ANOVA found no main effect of group (F (1, 580) = 0.939, p = 0.333).

The latDLPFC (Fig. 4C - left) however only exhibited significant signal increases in patients, and only during difficult cognitive task conditions (patients difficult onset: t = 10.129, p-FDR < 0.001; patients difficult block: t = 5.178, p-FDR < 0.001). Moreover, a two-way ANOVA found significant main effects of group (F (1, 580) = 20.38, p < 0.001), condition (F (3, 580) = 18.84, p < 0.001), and their interaction (F (3, 580) = 10.20, p < 0.001). Post hoc comparisons detected significantly greater latDLPFC activity in patients than controls at difficult onset (t = 7.003, p < 0.001).

Because latDLPFC activation was an exclusive characteristic of chronic pain-associated cognitive task networks, we investigated the possibility that latDLPFC function relates to pain. Dataset 1 obtained pain scans from healthy controls (n = 34) as mentioned previously, and patients (n = 105). FIR timeseries illustrated the absence of thermal stimulation-associated signal increases in medDLPFC (Fig. 4D - left) but their presence in latDLPFC (Fig. 4D - middle). One sample t-tests of activation values revealed significant increases in latDLPFC activity (Fig. 4D - right) in patients at the onset of both warm (t = 5.706, p-FDR < 0.001) and painful stimuli (t = 6.481, p-FDR < 0.001), as well as during the remainder of the pain block (t = 2.358, p-FDR = 0.041). Healthy controls also exhibited significant latDLPFC activity at pain onset (t = 2.265, p-FDR = 0.048) and during the pain block (t = 3.557, p-FDR = 0.003). Control group latDLPFC activation was less than patients (two-way ANOVA main effect of group: F (1, 548) = 11.38, p < 0.001, main effect of condition: F (3, 548) = 8.985, p < 0.001, interaction: F (3, 548) = 2.396, p = 0.067) during warm onset (post hoc: t = 3.08, p = 0.009) and pain onset (post hoc: t = 2.973, p = 0.012).

The medDLPFC, as part of both healthy control and migraine patient data-derived cognitive task networks, exhibited responses specific to cognitive task conditions in both groups. However, latDLPFC, which belonged exclusively to the cognitive task network of migraine patients, demonstrated pain-sensitive responses in patients and healthy controls. Thus, this pain-sensitive prefrontal cortex region displayed greater responses to pain in migraine patients than controls and was uniquely activated in patients during cognitive task performance. No patients reported experiencing a migraine or other pain during cognitive task scans.

Additionally, in Dataset 2, one sample t-tests detected significant increases in response to the auditory cue (Fig. 4E) in medDLPFC (t = 2.811, p-FDR = 0.013) and latDLPFC (t = 3.151, p-FDR = 0.008), but not to the subsequent onset (Fig. 4F, medDLPFC: t = 0.896, p-FDR = 0.470; latDLPFC: t = 0.216, p-FDR = 0.830) or offset of intense pain (Fig. 4G, medDLPFC: t = 1.025, p-FDR = 0.519; latDLPFC: t = 0.483, p-FDR = 0.632). This response profile mirrors that observed in LCL (auditory cue: t = 3.172, p-FDR = 0.008; intense onset: t = 1.034, p-FDR = 0.470; intense offset: t = 0.678, p-FDR = 0.627), while diverging from that of LaINS and RaINS (LaINS auditory cue: t = 1.929, p-FDR = 0.077; LaINS intense onset: t = 3.409, p-FDR = 0.008; LaINS intense offset: t = 3.926, p-FDR < 0.001; RaINS auditory cue: t = 1.437, p-FDR = 0.159; RaINS intense onset: t = 2.649, p-FDR = 0.029; RaINS intense offset: t = 5.003, p-FDR < 0.001).

### Migraine patients exhibit pathological claustrum activity

Because we hypothesize that the claustrum supports cognitive network initiation, we predicted that the altered cognitive task-associated network activity observed in patients was associated with altered claustrum activity. We therefore compared claustrum responses between patients and controls in cognitive task and pain conditions.

No significant claustrum activation group differences were observed during the difficult cognitive task (Fig. 5A-B). During pain, no group differences were present in LCL (Fig. 5C), but a two-way ANOVA revealed significant RCL activation differences between groups (Fig. 5D, main effect of group: F (1, 274) = 3.891, p = 0.0496; main effect of condition: F (1, 274) = 8.279, p = 0.004; interaction: F (1, 274) = 4.666, p = 0.032), with patients displaying greater RCL activity than controls at pain onset (post hoc: t = 2.922, p = 0.008). Claustrum effects were not attributable to subject movement (Supplemental Fig. S1B-C).

**Figure 5.**
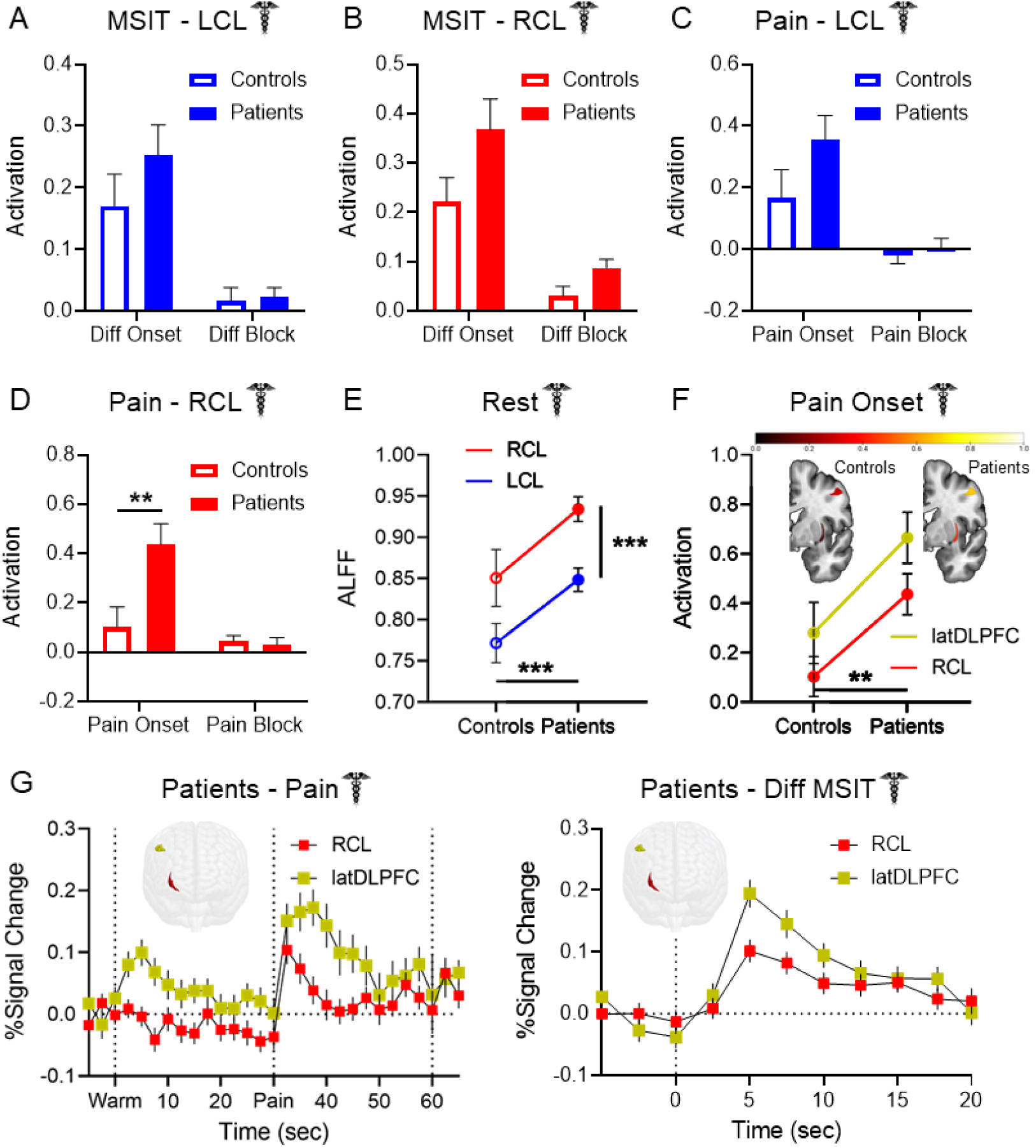
Pathological claustrum activity coincides with aberrant cognitive network region activity in migraine patients. No group differences were detected in **(A)** LCL or **(B)** RCL activation during cognitive task conditions between Dataset 1 healthy controls and migraine patients. **(C)** LCL activation during pain stimulation in Dataset 1 healthy controls and patients exhibited no significant group differences. **(D)** Patients exhibited significantly greater RCL activation than controls at pain onset (two-way ANOVA main effect of group: F (1, 274) = 3.891, p = 0.0496; condition: F (1, 274) = 8.279, p = 0.004; interaction: F (1, 274) = 4.666, p = 0.032; post hoc pain onset controls vs. patients: t = 2.922, p = 0.008). **(E)** Two-way ANOVA of ALFF detected significantly greater spontaneous claustrum activity at rest in migraine patients than controls (main effect of group: F (1, 280) = 13.30, p < 0.001), and in RCL than LCL (main effect of hemisphere: F (1, 280) = 14.07, p < 0.001). Post hoc comparisons found greater ALFF in patients than controls in RCL (t = 2.685, p = 0.015) and LCL (t = 2.473, p = 0.028). No significant interaction effect was observed (F (1, 280) = 0.022, p = 0.881). **(F)** Two-way ANOVA of RCL and latDLPFC activation at pain onset found significantly greater activation across regions in patients than controls (F (1, 274) = 8.503, p = 0.004). There was no significant main effect of region (F (1, 274) = 2.699, p = 0.102) or significant interaction (F (1, 274) = 0.044, p = 0.834). Heat maps of RCL (bottom structure) and latDLPFC (top structure) similarly depicted greater activation at pain onset across regions in patients (right) than controls (left). **(G)** FIR timeseries illustrated coincident increases in RCL and latDLPFC BOLD percent signal change in Dataset 1 patients at **left:** the onset of pain and **right:** the onset of the difficult cognitive task. Timeseries are consistent with changes at condition onsets due to hemodynamic delay. No statistical tests were performed on these timeseries. 3-D rendered brains display shape and location of RCL (red) and latDLPFC (gold).

Patients also displayed greater bilateral anterior insula activity at pain onset than controls (Supplemental Fig. S2). Although post hoc comparisons found no group differences in either ROI specifically, a two-way ANOVA detected a main effect of group (F (1, 274) = 7.269, p = 0.007) across LaINS and RaINS.

It was previously reported that migraine patients exhibit increased Amplitude of Low Frequency Fluctuations (ALFF) than controls in RCL at rest (Gu et al., 2023). ALFF is a putative measure of spontaneous neuronal activity obtained via fMRI (Zang et al., 2007), and the convergence of this finding with the RCL group differences detected in our study compelled us to attempt a replication of the ALFF effect. Dataset 1 acquired resting state BOLD scans from healthy controls (n = 33) and patients (n = 109). A two-way ANOVA comparing LCL and RCL resting state ALFF between groups (Fig. 5E) identified main effects of group (F (1, 280) = 13.30, p < 0.001) and hemisphere (F (1, 280) = 14.07, p < 0.001) but not of their interaction (F (1, 280) = 0.022, p = 0.881), such that patients exhibited greater ALFF than controls in both regions and RCL exhibited greater ALFF than LCL in both groups. Post hoc comparisons affirmed that ALFF was greater in patients than controls in RCL (t = 2.685, p = 0.015) and LCL (t = 2.473, p = 0.028). However, no significant differences were present when measuring fractional ALFF (fALFF; Supplemental Fig. S3), a measure of ALFF as a proportion of the amplitudes of all frequencies obtained (Zou et al., 2008). No group differences were found in latDLPFC, medDLPFC, DMN, or Extrinsic Mode Network (EMN) resting state ALFF.

Whole-brain seed-to-voxel functional connectivity of LCL, RCL, medDLPFC, and latDLPFC seeds were also compared between Dataset 1 controls and patients. A group difference was only identified in latDLPFC, with patients exhibiting decreased latDLPFC functional connectivity with a cluster comprising voxels in the right orbitofrontal cortex and inferior frontal gyrus (peak: (54, 32, -10); size: 124; family-wise error rate corrected p-FWE: 0.042).

### Right claustrum and latDLPFC co-activate across multiple experimental conditions

The claustrum is hypothesized to project to cognitive control regions to support network initiation, and latDLPFC is a probable ipsilateral RCL projection target (Markowitsch et al., 1984; Reser et al., 2014; Tanné-Gariépy et al., 2002; Torgerson et al., 2015; Wang et al., 2023; White et al., 2017). Therefore, upon identifying differences between patients and controls in both regions, we investigated the possible link between RCL and latDLPFC activity. A two-way ANOVA comparing RCL and latDLPFC activation in Dataset 1 at pain onset between healthy controls and patients (Fig. 5F) detected a significant main effect of group (F (1, 274) = 8.503, p = 0.004), with signal in patients greater than in controls across regions. FIR timeseries of Dataset 1 RCL and latDLPFC signal also illustrated contemporaneous increases in patients at the onset of pain (Fig. 5G - left) and the onset of the difficult cognitive task (Fig. 5G - right).

### Structural connectivity in RCL-latDLPFC and RCL-medDLPFC circuits in healthy individuals

After observing the co-activations reported above, we sought to verify the presence of anatomical connections between RCL our DLPFC ROIs. High resolution (7T) DWI scans were analyzed in a sample of healthy individuals (n = 174) from the Human Connectome Project (HCP; Van Essen et al., 2013). Tractography analyses were run to assess white matter connectivity in the hypothesized RCL-latDLPFC and RCL-medDLPFC circuits.

White matter connectivity was identified for RCL-latDLPFC (W = 15225, p-FDR < 0.001) and RCL-medDLPFC (W = 15225, p-FDR < 0.001), and a direct comparison found a significantly stronger RCL-medDLPFC connection (Fig. 6A, W = 12607, p-FDR < 0.001). This indicates preferential structural connectivity between RCL and medDLPFC, over latDLPFC, in healthy individuals. Tractogram analyses show that white matter connections between RCL and either latDLPFC or medDLPFC take two paths: (1) traveling from RCL posteriorly and superiorly along the middle longitudinal fasciculus to the arcuate fasciculus before joining the superior longitudinal fasciculus and on to the medDLPFC/latDLPFC; or (2) traveling from RCL directly superior along the superior thalamic radiation before connecting to the superior longitudinal fasciculus and then to the medDLPFC/latDLPFC. RCL-latDLPFC favors path 2, but the circuits appear to overlap superior to the insula before the RCL-latDLPFC circuit branches along the superior longitudinal fasciculus to latDLPFC (Fig. 6A).

**Figure 6.**
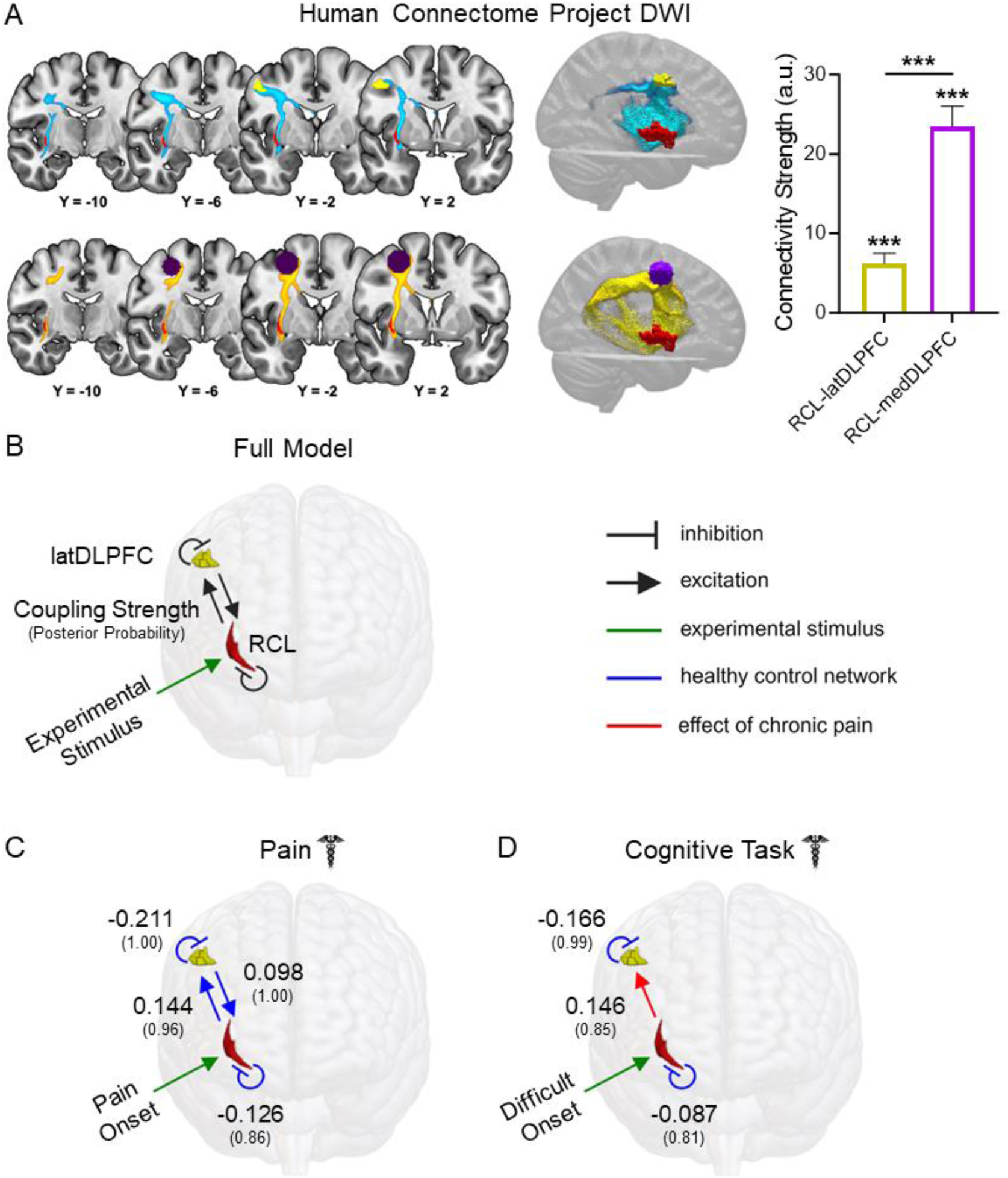
Structural and effective connectivity are consistent with an excitatory RCL→latDLPFC projection altered in migraine patients. **(A)** Group tractograms of **(left-top)** RCL-latDLPFC and **(left-bottom)** RCL-medDLPFC structural connectivity thresholded at 50% (indicating WM fibers detected in at least 50% of individuals sampled) in healthy HCP participants (n = 174). **Right:** Median structural connectivity strength (a.u.) of RCL-latDLPFC and RCL-medDLPFC circuits in healthy HCP participants with 95% CI displayed. Both circuits showed significant structural connectivity (RCL-latDLPFC: W = 15225, p-FDR < 0.001; RCL-medDLPFC: W = 15225, p-FDR < 0.001), and RCL-medDLPFC showed significantly greater connectivity strength than RCL-latDLPFC (W = 12607, p-FDR < 0.001). **(B)** Fully connected DCM models with bidirectional RCL←→latDLPFC excitatory projections and self-inhibitory connections in both ROIs were modeled for Dataset 1 pain and cognitive task scans separately. Experimental stimuli (green arrow) were modeled to affect RCL due to the a priori circuit hypothesis. Here, modeled connections are displayed in black. In subsequent panels, blue lines represent connections detected on average across healthy participants. Red lines depict effects of chronic pain and therefore represent significant group differences. Large font numbers represent second level coupling parameters (positive = excitatory), and small font numbers in parentheses represent posterior probabilities quantifying the strength of evidence for each coupling parameter (0.00 – 1.00). A posterior probability exceeding 0.75 is considered “positive evidence.” Therefore, only parameters with posterior probability greater than 0.75 were included in figure panels. **(C)** DCM found evidence of bidirectional excitatory RCL←→latDLPFC connectivity and self-inhibition in both ROIs at pain onset in healthy controls. No effects of chronic pain were detected for this model, meaning effective connectivity in patients at pain onset is not significantly different than controls. **(D)** At difficult cognitive task onset, DCM only found evidence of self-inhibition in RCL and latDLPFC in healthy controls, with no evidence for excitatory projections between regions. However, evidence was found for increased RCL→latDLPFC effective connectivity due to chronic pain, consistent with the appearance of an excitatory projection during cognitive task processing in patients.

### Dynamic causal modeling suggests an excitatory RCL**→**latDLPFC projection altered in migraine patients

In addition to verifying RCL-latDLPFC anatomical connectivity, we performed dynamic causal modeling (DCM) to see if the regions’ BOLD timeseries were consistent with a possible RCL→latDLPFC projection. Unlike functional connectivity analyses, which are correlational, DCM (Friston et al., 2003) uses a Bayesian framework to assess the strength of evidence for a directional effect of one region on another. For every connection modeled, values are obtained describing coupling strength (positive for excitatory connections) and the strength of evidence (posterior probability). We only report results exceeding the threshold for “positive” evidence (greater than 0.75 posterior probability).

Fully connected models comprising bidirectional projections between RCL and latDLPFC as well as inhibitory self-connections in each ROI (Fig. 6B) were estimated for all included subjects in pain and cognitive task scans separately (Zeidman et al., 2019a). Group level analyses were conducted by specifying a fully connected Parametric Empirical Bayes (PEB; Zeidman et al., 2019b) model and searching through all possible reduced models. Because RCL is hypothesized to route information to latDLPFC, all models specified effects of experimental input (i.e., pain onset, difficult cognitive task onset) on RCL. To compare effective connectivity between patients and controls, two second level PEB regressors were coded to model: 1) average network parameters among healthy controls (blue lines in figures) and 2) the change in network parameters due to chronic pain (i.e., group differences – red lines in figures). At acute pain onset (Fig. 6C), evidence was found in healthy controls for inhibitory self-connections in RCL (coupling strength: -0.126, posterior probability: 0.86) and latDLPFC (-0.211, 1.00), as well as bidirectional excitatory projections (RCL→latDLPFC: 0.144, 0.96; latDLPFC→RCL: 0.098, 1.00). No network changes due to chronic pain exceeding 0.75 posterior probability were detected, meaning effective connectivity in patients at pain onset was not significantly different than controls. At difficult task onset (Fig. 6D), evidence was found in healthy controls only for inhibitory self-connections (RCL: -0.087, 0.81; latDLPFC: -0.166, 0.99), but not for excitatory projections between regions. However, chronic pain was found to increase RCL→latDLPFC effective connectivity (0.146, 0.85), consistent with the appearance of an excitatory projection during cognitive task processing in patients.

## Discussion

In this study, we analyzed human fMRI data to investigate the relationship between claustrum and cognitive control network activity during acute and chronic pain. Our results indicate the human claustrum responds to acute thermal pain and pain cues. Increased RCL activity at the onset of acute pain was observed in migraine patients. This pathological claustrum activity accompanied altered cognitive task-associated network activity in patients, which was characterized by the additional recruitment of a pain-responsive cognitive control network region (latDLPFC), in the absence of acute pain. DWI verified RCL-latDLPFC structural connectivity, and DCM findings were consistent with an excitatory RCL→latDLPFC projection during acute pain in healthy participants, as well as the appearance of RCL→latDLPFC effective connectivity during cognitive task processing in patients with chronic pain.

### The human claustrum is responsive to pain or pain-predictive cues

The claustrum exhibited responses in healthy participants to acute thermal pain, a stimulus that induces activation in cognitive task-associated network regions (Seminowicz & Davis, 2007). In a separate sample of healthy participants where pain was preceded by an auditory cue, significant claustrum activation was associated with pain anticipation, rather than pain onset. While claustrum activation at pain onset would be predicted by a salience processing hypothesis of claustrum function (Remedios et al., 2014), the absence of significant claustrum responses to the onset of pain, an inherently salient stimulus, after a cue in Dataset 2 challenges this interpretation. Additionally, while a salience-directed attention claustrum function (Smith et al. 2019) may still account for these observations, the remainder of our findings compel us to favor the Network Instantiation in Cognitive Control (NICC) model (Madden et al., 2022), which proposes that the claustrum supports cortical network formation for cognitive control.

We observed structural connectivity between claustrum and DLPFC regions (medDLPFC & latDLPFC) as well as their coincident activation in multiple conditions, including analogous shifts in activation from pain onset to pain cue between experiments in claustrum and latDLPFC. Indeed, claustrum responses across experiments aligned with regions of DLPFC, which is robustly associated with executive functioning. Importantly, conditions were identified (e.g., pain onset when preceded by a cue, pain offset) where claustrum responses diverged from the anterior insula, a primary region of the salience network.

The claustrum’s association with executive function is additionally supported by the prior finding that the claustrum displays widespread resting state functional connectivity with cognitive control network regions (Krimmel et al., 2019) and this study’s DCM evidence of an excitatory RCL→latDLPFC projection in response to pain onset in healthy controls. Remarkably, latDLPFC is not part of the Neurologic Pain Signature (Wager et al., 2013), a whole-brain fMRI pattern predictive of reported pain intensity. This implies latDLPFC activity, supported by RCL, may encode cognitive rather than nociceptive components of pain, such as the effect of prior pain experience (Melzack & Casey, 1968).

### Migraine patients engage a pain-responsive prefrontal cortex region during pain-free cognitive task processing

Our finding of elevated cognitive task-associated activity across multiple clusters in migraine patients echoes previous studies identifying greater task-induced activity (increased activation/decreased deactivation) in chronic pain patients than controls (Baliki et al., 2008; Ceko et al., 2015; Mathur et al., 2015; Seminowicz et al., 2011). However, to the best of our knowledge this study is the first to empirically characterize the function of a region exhibiting task-induced group differences. Our decision to analyze a particular right DLPFC region exhibiting greater cognitive task-associated activity in patients than controls was motivated by its prominence in the group differences report and its relationship to the primary cognitive task LV of healthy participants. Specifically, the chosen cluster was the largest and most statistically significant region displaying group differences, and it contained a substantial ROI (latDLPFC) outside the healthy control cognitive task network. The revelation that latDLPFC, an additional cognitive task network region in migraine patients, is related to processing pain and pain cues in healthy conditions invites deeper exploration into the transition to, and potentially the subjective experience of, chronic pain.

The DLPFC is large and functionally heterogeneous. Comparisons with other primates reveal that the human DLPFC experienced considerable evolutionary expansion (Van Essen & Dierker, 2007). It is consequently believed to participate in higher order cognitive processing, and this is supported by experiments implicating DLPFC in numerous functions including attention (Bidet-Caulet et al., 2014), working memory (Barbey et al., 2013), decision-making (Rahnev et al, 2016), and emotional regulation (Buhle et al., 2014). Notably, studies have associated DLPFC activity with cognitive components of pain such as expectations of pain (Atlas et al., 2010), placebo analgesia (Krummenacher et al., 2010), pain tolerance (Graff-Guerrero et al., 2005; Lorenz & Casey, 2003), perceived control of pain (Wiech et al., 2006), and pain catastrophizing (Seminowicz & Davis, 2006), including in migraine patients (Hubbard et al., 2014). Mindfulness meditation, which is intended to help practitioners reframe their pain experience, is also associated with modulation of DLPFC activity (Allen et al., 2012).

Intriguingly, fMRI investigations of non-nociceptive aspects of pain have reported activation in coordinates near latDLPFC peak activity. These include tests of “virtual” pain in which healthy participants viewed a video of a needle puncturing a palm (Ushida et al., 2008) and an experiment where migraine patients observed pain-related words during a state of distracted attention (Eck et al., 2011). A study of simultaneous cognitive task performance and capsaicin-induced hyperalgesia in healthy participants also reported activation coordinates near latDLPFC associated with the interaction of pain intensity and cognitive demand (Wiech et al., 2005). However, these studies did not compare chronic pain patients and healthy controls during a difficult cognitive task, and they did not acquire pain and cognitive task scans from both groups. They therefore were unable to identify latDLPFC as an additional pain-responsive cognitive network region in chronic pain. When considering these results with our findings of increased latDLPFC activity in chronic pain patients at the onsets of both acute pain and a difficult cognitive task, it is possible that latDLPFC activity is associated with the cognitive load of pain. For example, increased latDLPFC activity among migraine patients may be associated with an increased propensity to factor pain, potential pain, or prior pain into decision making processes.

Although medDLPFC and latDLPFC responses in healthy controls were specific to cognitive task and pain processing respectively in Dataset 1 of our study, this is likely due to the absence of other conditions. The variety of conditions associated with DLPFC activity, as well as the dual medDLPFC and latDLPFC responses to a pain-predictive cue in our Dataset 2 findings, hint at more domain-general roles for both regions. Considerable evidence describes DLPFC functional gradients along several axes (Abdallah et al., 2022; Jung et al., 2022; Koechlin et al., 2003; Nee & D’Esposito, 2016; Petrides, 2005), so it is reasonable to predict medDLPFC and latDLPFC inhabit different positions along a functional continuum rather than perform fundamentally distinct roles. Ultimately, the divergence of medDLPFC and latDLPFC activity between conditions and groups observed in this study merits further investigation to determine the generalizability of our finding, to test the relationship between latDLPFC activity and symptoms of chronic pain, such as executive function impairments, and to probe the mechanisms producing chronic pain-associated changes in latDLPFC activity.

### Pathological claustrum activation drives aberrant cognitive network processing in chronic pain

Our study provides the first clues that claustrum dysfunction may underlie latDLPFC and other cognitive network changes in chronic pain. Because we hypothesized that such changes result from altered claustrum activity upstream of cognitive control network initiation, we tested for evidence of an excitatory RCL→latDLPFC projection that is altered in migraine patients. RCL and latDLPFC suggestively co-activated in multiple conditions, including during difficult cognitive task onset in patients. High-resolution DWI verified RCL-latDLPFC structural connectivity, and effective connectivity analyses pointed to a possible causal influence of RCL on latDLPFC activity. DCM of Dataset 1 pain scans found evidence of bidirectional RCL←→latDLPFC excitatory projections at pain onset in healthy controls, with no significant changes due to chronic pain. Strikingly, DCM of Dataset 1 cognitive task scans found no evidence of RCL←→latDLPFC coupling in controls at difficult task onset, but evidence was found of increased excitatory RCL→latDLPFC effective connectivity in patients. Our findings are therefore consistent with an RCL→latDLPFC projection associated with processing cognitive components of pain in healthy conditions that is aberrantly recruited during cognitive task processing in chronic pain due to underlying claustrum dysfunction.

Recent studies of rodent chronic pain models also implicate the claustrum in chronic pain, and their findings are consistent with a link between the claustrum and chronic pain-associated cognitive network changes. Specifically, in different chronic pain models Xu et al. (2022) and Ntamati et al. (2023) identified changes in circuitry between the claustrum and anterior cingulate cortex, a primary node of cognitive control brain networks (Menon & D’Esposito, 2022). In humans, Gracely et al. (2004) observed positive correlations in fibromyalgia patients between claustrum activity and pain catastrophizing, a hallmark cognitive component of pain.

These findings raise the possibility of the claustrum as a therapeutic target in chronic pain. In particular, psychedelics are hypothesized to exert therapeutic effects by reorganizing activity and connectivity among large scale brain networks (Carhart-Harris & Friston, 2019; Castellanos et al., 2020; Doss et al., 2022; Vollenweider & Geyer, 2001), and psilocybin alters claustrum functional connectivity with cortical networks (Barrett et al., 2020). Therefore, our findings and recent evidence indicating reduced migraine frequency after psilocybin (Schindler et al., 2021) position the claustrum as a potential driver of benefits from psychedelic-assisted therapy for chronic pain.

## Limitations

Despite testing the claustrum and other ROIs in multiple conditions and experimental paradigms, this study contained no direct brain manipulation. Although our findings are consistent with the NICC model, in which the claustrum drives activity in cognitive control network regions, the causal relationship between the claustrum and other analyzed ROIs, such as latDLPFC, is undetermined. Furthermore, this study cannot determine if increased RCL BOLD at pain onset in patients arises due to changes within RCL or changes elsewhere in the system, such as sensory or association cortices upstream.

It also remains to be seen if the group differences detected between healthy controls and migraine patients in RCL and latDLPFC activity are present in other chronic pain conditions. Notably, our migraine patient cohort was predominantly female, as was our Dataset 1 sample of healthy controls. It is therefore possible the generalizability of the group differences is limited by untested sex effects.

Lastly, in Dataset 1, LCL exhibited significant signal increases at pain onset as predicted, and RCL had generally greater signal during pain than warmth across timepoints. This laterality difference could result from the thermal stimulus application to the left side of the body. However, additional laterality effects, such as unilateral LCL responses to a pain-predictive cue, detected in this study and others (Barrett et al., 2020; Gu et al., 2023) are consistent with the possibility of laterality in claustrum function. Unfortunately, uniform application of thermal stimuli to the left side of the body in this study rendered us unable to assess laterality effects. Furthermore, the absence of patients with chronic pain in our Dataset 2 sample prevented us from testing if RCL also exhibits increased responses to pain cues in patients compared to controls.

## Conclusions

Using neuroimaging data from healthy participants and migraine patients, our study reveals a relationship between claustrum and cognitive control network activity during acute pain and uncovers evidence of a claustrum-DLPFC circuit underlying cognitive network dysfunction in chronic pain. These findings represent empirical support for the Network Instantiation in Cognitive Control model of claustrum function and raise the possibility of the claustrum as a future therapeutic target for chronic pain conditions, with potential implications for other neuropsychiatric disorders characterized by cognitive impairment and cortical network abnormalities.

## Acknowledgments

We thank Dr. AR McIntosh for supporting our use of PLS Software. We thank Dr. AJ Shackman for sharing the negative affect, pain, and cognitive control meta-analysis database. We thank ZS Sidhu for assistance in analyzing claustrum responses to a pain-predictive cue. This work was supported by R01 NS112356, R01 AT007176 to DAS.

## Materials and methods

### Participants

fMRI analyses were performed in two datasets. Dataset 1 was acquired from 112 (98 female, mean age = 36.79) individuals with at least a one-year history of migraine diagnosis (mean headache days/month = 9.0) and 35 healthy subjects (30 female, mean age = 37.44) recruited as controls in a clinical trial at the University of Maryland, Baltimore Medical Imaging Facility. Data for resting-state (patients n = 109; controls n = 33), cognitive task (patients n = 112; controls n = 35), and pain (patients n = 105; controls n = 34) scans represent either the entirety or subsets of the overall dataset depending on scan acquisition and quality control parameters, such as subject motion. For full details of clinical trial recruitment, see Seminowicz et al. (2020). The clinical trial performed three longitudinal imaging sessions, one at baseline and two after the onset of therapeutic intervention. Only baseline scans were analyzed for patients. No patient participants used medication within 24 hours of a scanning session or experienced a headache during image acquisition.

Because healthy controls did not undergo therapeutic intervention and analyses demonstrated no significant changes in variables of interest over time in this group, control participant data from all sessions were averaged within subjects.

Dataset 2 acquired data from 49 healthy participants. Exclusion of participants due to incidental radiological findings (n = 5), hardware malfunctions (n = 2), participant noncompliance (n = 2), and reports of pain not attributable to experimental stimuli (n = 1), yielded a final sample n = 39 (21 female, mean age = 27.79).

For both datasets, data collection was approved by the University of Maryland, Baltimore Institutional Review Board, and informed written consent was obtained from each participant prior to any study procedures.

### fMRI Acquisition and Preprocessing

Dataset 1 images were acquired with a Siemens 3T Tim Trio scanner with a 32 channel head coil (patients n = 112, controls n = 21) or a Siemens 3T Prisma scanner with a 64 channel head coil (controls n = 13) due to a scanner upgrade during acquisition. For one control participant, the initial scan session was performed with the 3T Trio pre-upgrade, and subsequent sessions were performed with the 3T Prisma post-upgrade. Each scanning session acquired a T1 MPRAGE (repetition time [TR] 2300 ms, echo time [TE] 2.98 ms, slice thickness 1mm, field of view [FOV] 256mm, flip angle 9°, and voxel size 1 x 1 x 1 mm) high-resolution anatomical scan for template registration, an fMRI scan with blocks of thermal stimulation (two runs of eight minutes, echo planar imaging [EPI], TR 2500 ms, TE 30 ms, slice thickness 3 mm, FOV 230 mm, flip angle 90°, and voxel size 3 x 3 x 3 mm), an fMRI scan with blocks of cognitive task stimuli (two runs of five minutes, EPI, TR 2500 ms, TE 30 ms, slice thickness 3 mm, FOV 230 mm, flip angle 90°, and voxel size 3 x 3 x 3 mm), and a resting state fMRI scan (one run of 10 minutes, EPI, TR 2000 ms, TE 30 ms, slice thickness 3 mm, FOV 230 mm, flip angle 90°, and voxel size 3 x 3 x 3 mm).

All Dataset 1 images were preprocessed in SPM12 (http://www.fil.ion.ucl.ac.uk/spm/software/spm12/). Preprocessing included slice timing correction, realignment (motion correction), coregistration of T1-weighted structural scans to mean realigned functional images, segmentation of structural scans, and normalization of structural and realigned functional images to a standard MNI template with interpolation to 2 x 2 x 2 mm voxels using a 4th degree B-Spline. For analyses of whole-brain cognitive task processing (i.e., PLS) and resting state functional connectivity, images were smoothed with a 6mm full width at half maximum (FWHM) Gaussian kernel. In all other analyses, the data was not smoothed to prevent inclusion of extra-claustral signal in claustrum ROI voxels. Claustrum functional connectivity was similarly calculated using unsmoothed mean LCL and RCL timeseries.

Dataset 2 images were acquired with a Siemens 3T Prisma scanner with a 64 channel head coil. Each scanning session acquired a T1 MPRAGE (TR 2300 ms, TE 2.94 ms, slice thickness 1 mm, FOV 256 mm, flip angle 9°, and voxel size 1 x 1 x 1 mm) high-resolution anatomical scan for template registration and up to five fMRI scans with blocks of thermal stimulation (8 minutes, EPI, TR 1750 ms, TE 35 ms, slick thickness 3 mm, FOV 240 mm, 75° flip angle, and voxel size 3 x 3 x 3 mm).

All Dataset 2 images were preprocessed in SPM12 (http://www.fil.ion.ucl.ac.uk/spm/software/spm12/). Preprocessing included realignment (motion correction), coregistration of T1-weighted structural scans to mean realigned functional images, segmentation of structural scans, and normalization of structural and realigned functional images to a standard MNI template with interpolation to 2 x 2 x 2 mm voxels using a 4th degree B-Spline. To prevent inclusion of extra-claustral signal in claustrum ROI voxels, the data was not smoothed.

### Tasks

In Dataset 1 pain scans, a moderately painful (5-7 on a 0-10 numeric rating scale) temperature was selected for each subject based on responses to pre-scan quantitative sensory testing and confirmed with verbal ratings when subjects were inside the scanner. Thermal stimuli were applied to the left forearm using a Medoc Inc Pathway with ATS 30 mm x 30 mm thermode. Each run included five blocks of non-noxious warm stimulation (two second ramp up from 32°C and 28 second hold) defined as a temperature 8°C less than the painful temperature. Warm stimulation was immediately followed by the subject-specific painful stimulation (two second ramp up and 28 second hold) and a two second descending ramp to 32°C (baseline). Intertrial intervals consisted of a 28 second hold at 32°C (baseline).

For Dataset 1 cognitive task scans, participants performed the multi-source interference task (MSIT; Bush et al., 2003). The task was administered as previously reported (Seminowicz and Davis, 2007; Seminowicz et al., 2011). Briefly, in each trial an array of three numbers was presented in which two numbers were identical and one number differed, and the participant was directed to press one of three buttons on a response pad corresponding to the unique number. There were two difficulty levels (easy, difficult), which were performed in separate 20s blocks (10 trials per block). In the easy condition, the unique number matched its position in the array (e.g., “1-3-3”, “1-2-1”, “2-2-3”). In the difficult condition, the unique number did not match its position in the array (e.g., “3-1-1”, “2-1-2”, “3-3-2”). A sequential tapping task was used as a control condition in which an asterisk appeared on the screen in one of three positions and the participant pressed a button corresponding to the position of the asterisk. Subjects were trained on the MSIT outside of the scanner before performing the task during fMRI scans.

For Dataset 2 scans, thermal stimulation was delivered at multiple intensities and durations in a pseudorandom order. As in Dataset 1, a temperature rated as inducing a 50 pain level on a 0-100 rating scale was selected for each subject based on responses to pre-scan quantitative sensory testing and confirmed with verbal ratings when subjects were inside the scanner. This temperature was defined as “moderate” pain, and additional “intense” and “slight” pain levels were defined as 1°C greater and 1°C less than the moderate pain temperature respectively. A control condition of non-painful warm stimulation was uniformly defined as 38°C across subjects. After an inter-trial interval of 16 or 20 seconds, each trial began with a doorbell sound audio cue, which was followed by an anticipation period of 7.5 seconds prior to thermal stimulation. Thermal stimuli were applied to the left lower leg using a Medoc Inc Pathway with CHEPS (27 mm diameter) thermode. Stimulation began with a two second ramp up from baseline temperature (32°C) to the trial’s target temperature, and stimulation lasted 4, 8, 21, or 36 seconds (pseudorandom) before a two second ramp down to baseline temperature. The ramp down was immediately followed by a prompt for the participant to rate their pain experience in that trial on a 0-100 scale in intervals of 10 using a response pad. Each run consisted of 10 trials, and as many as five runs were obtained from each participant.

For all task-based analyses, functional scans exhibiting an average Framewise Displacement greater than 1.0 mm were excluded for excessive motion.

### PLS Analyses of Cognitive Task Network Activity

PLS software (McIntosh et al. 1996) was used to identify whole-brain network activity associated with increased cognitive task difficulty from smoothed (6mm FWHM) Dataset 1 cognitive task scans. In a similar manner to principal component analysis, PLS yields “latent variables” (LVs) representing multivariate patterns of brain activity covarying with experimental conditions and ordered by the percent of variance they explain. To assess LV significance, 500 permutation tests were performed in which condition labels (i.e., “easy”, “difficult”, “tapping”) were shuffled among condition blocks and results were compared to the unshuffled design, yielding a p-value for each LV. Multiple comparisons correction is not necessary in these analyses because permutations assess the significance of entire multivariate patterns rather than of individual voxels. The reliability of each voxel within an LV was assessed with 100 bootstrap samples in which subject data was resampled with replacement within condition and the PLS was run with each sample set. This yields a bootstrap ratio (BSR) for each voxel within the LV analogous to a z-score, where a BSR of 1.96 equals an uncorrected p-value of 0.05. LVs were thresholded at BSR +/- 3 which equates to an uncorrected p-value of 0.0027 and is standard for PLS analysis.

Within group analyses of cognitive task network activity were conducted with the software default mean-centering option, namely removing group means from condition means to emphasize condition effects.

The patients versus controls comparison was conducted after removing grand condition means from group condition means to assess how group membership modulates conditions. The cluster report from this group effects analysis included clusters of at least 50 voxels at least 10mm apart.

### Regions of Interest

LCL, RCL, insular cortex, and putamen ROIs were drawn by hand on the normalized anatomical images of 20 subjects in a publicly available dataset (n = 22) acquired on a 7T MR Scanner (MAGNETOM 7T, Siemens Healthcare, Erlangen, Germany), and a group average ROI file was obtained for each region. Further details of the scans can be found in Gorgolewski et al. (2015). Two subjects were omitted from ROI generation, one due to preprocessing errors preventing normalization of sufficient quality and one due to acquisition via different scanning parameters.

LaINS and RaINS ROIs were generated in MarsBaR (Brett et al., 2002) based on centers of mass and volumes derived from a coordinate based meta-analysis of imaging studies of cognitive control, pain, and negative affect (https://identifiers.org/neurovault.collection:474; Shackman et al., 2011). Spherical ROIs were generated with the following parameters: LaINS (center: -36, 13, 2; radius: 8mm), RaINS (center: 36, 16, 2; radius: 8mm).

latDLPFC was defined based on PLS analysis of group differences during cognitive task conditions (see above). The latDLPFC region was derived from the primary cluster exhibiting group differences, with a peak voxel at 52, 0, 44. Binarized PLS results were masked with a cube positioned to encompass only voxels within this cluster that additionally did not overlap with any voxels contained within the healthy control cognitive task primary LV. This region was then masked with the second level pain scan mask from healthy controls to ensure the latDLPFC ROI only included voxels where signal was obtained from both groups.

medDLPFC was also defined based on PLS analyses. A 10mm sphere was generated with a center at 28, 0, 52 because it fell within the primary cognitive task associated LV for both healthy controls and patients.

DMN and EMN masks were also derived from PLS analyses. The primary LV of cognitive task associated activity in healthy controls was binarized and thresholded at either side of 0 to yield binarized DMN and EMN ROIs.

Flanking region ROI definition is explained below.

### Small Region Confound Correction

As described in Krimmel et al. (2019), the effect of neighboring insular cortex and putamen on claustrum signal in resting state data was controlled via SRCC. Insular cortex and putamen “flanking” ROIs were defined by dilating each hemisphere’s claustrum ROI four functional voxels and identifying the overlap between the dilated claustrum ROIs and their neighboring insula and putamen ROIs at least two functional voxels separated from the original claustrum. This generated “flanking” ROIs within the insular cortex and putamen, similar to the claustrum’s shape, yet distant from the claustrum to avoid including claustrum signal. Timeseries from these regions were included as additional regressors to control for their effects in resting state claustrum analyses.

An additional step was added to this protocol to control for insular cortex and putamen signal in task-based claustrum analyses. If flanking ROI signals are influenced by task conditions, using raw timeseries from flanking ROIs as additional regressors risks removing condition-induced variation from the claustrum signal. Therefore, when analyzing claustrum signal in task scans, the CONN Toolbox (RRID:SCR_009550; Nieto-Castanon, 2020) was used to generate canonical-HRF-convolved condition timeseries, and a regressor was generated for each flanking ROI by obtaining the interaction of the ROI’s raw timeseries and the summed HRF-convolved timeseries of all modeled conditions. This process yielded a timeseries for each flanking ROI that covaried with the ROI physiological timeseries during task conditions but lacked any variation potentially induced by the conditions, allowing control of the influence of neighboring regions on claustrum signal without indirectly removing task effects.

### Finite Impulse Response Models

To analyze claustrum responses to conditions over time, and because the claustrum is not a laminar cortical structure, a GLM was fitted using an FIR to model the HRF. This yielded task-associated claustrum signal change timeseries without assuming the dynamics of the claustrum’s HRF.

For Dataset 1 experimental pain analyses, a single “stimulation” condition was modeled encompassing the entire non-noxious warmth and painful stimulation blocks, as well as five seconds of pre-warm baseline stimulation and five seconds of post-pain temperature ramp down and baseline stimulation. This allowed extraction, via MarsBaR software, of percent signal change FIR timeseries encompassing pre-stimulus baseline stimulation, non-noxious warm stimulation, pain stimulation, and pain offset timepoints against an implicit baseline comprised of inter-trial interval timepoints.

For Dataset 1 cognitive task analyses, the easy and difficult task levels were separately modeled as “easy” and “difficult” conditions that encompassed the entire task block as well as two pre-stimulus timepoints, allowing extraction in MarsBaR of percent signal change FIR timeseries against an implicit baseline comprised of tapping control condition timepoints.

For all analyses, trial timeseries were averaged within subjects and then within groups. Treating trial onset as t0, averaged percent signal change timeseries were transformed by averaging pre-stimulus baseline TRs (t-2 & t-1) across the group and subtracting this value from every time point. This allowed comparisons of post-stimulus-onset signal values to pre-stimulus values while accounting for variability in pre-stimulus baseline signal.

### Task-Induced Activation Models

To make direct comparisons between conditions, regions, and groups of task-induced activation, singular activation values were derived from GLM analyses in SPM12. Due to the a priori hypothesis that claustrum responds transiently to stimulus onset, all conditions were modeled as separate “onset” and “block” conditions. Because Dataset 1 pain scans used a two second temperature ramp up, to keep analyses uniform, “onset” conditions were always defined as the first two seconds of a condition, and “block” conditions were always defined as the remainder of the condition.

Dataset 2 stimulation trials also ramped up over two seconds to target temperatures, but painful trials began at a 32°C baseline instead of a higher non-painful warm temperature as in Dataset 1. This resulted in the absence of significant signal increases in salience network regions during the ramp up condition regardless of target temperature. Therefore, Dataset 2 models included an “upramp” condition consisting of the two second ramp up as well as an “onset” condition consisting of the first two seconds of stimulation at the trial’s target temperature prior to “block” and “offset” conditions.

For analyses of ROIs other than the claustrum, GLMs included all task conditions and six motion parameters. Second level results were separately masked with each ROI and activation values were averaged across the ROI within each subject, and then across subjects within each group. Separate GLMs were used to analyze each claustrum ROI, which included all task conditions, six motion parameters, and the two flanking region-condition interaction timeseries generated via the task-adapted SRCC process described above.

### Resting State Functional Connectivity

Resting state scans were acquired while participants fixated on a plus sign. Scans exhibiting an average Framewise Displacement greater than 0.5 mm were excluded for excessive motion. This resulted in the exclusion of eight resting state scans. Three were from patient baseline scans, resulting in removal of three patient participants. Five were individual runs from five different healthy participants, resulting in the removal of two healthy controls from whom only one resting state scan was acquired.

Analyses of preprocessed resting state functional images were performed in the CONN Toolbox (RRID:SCR_009550; Nieto-Castanon, 2020). Nuisance regressors included motion parameters and their first order derivatives, a scrubbing vector generated by ART-toolbox identification of outlier scans (global-signal z-value threshold = 5; subject-motion mm threshold = 0.9), and the first five principal components of white matter (WM) and cerebrospinal fluid (CSF) masks (aCompCor; Behzadi et al., 2007; Muschelli et al., 2014). Because healthy controls and migraine patients were hypothesized to exhibit different network phenotypes, global signal regression was not performed to avoid differential group impacts (Murphy and Fox, 2017). Consequently, a 2x eroded CSF mask and a 4x eroded WM mask, not containing external or extreme capsules, were used because such masks no longer contain global signal (Power et al., 2017). Regression was performed simultaneously with band-pass filtering (Hallquist et al., 2013) between 0.008 Hz and our acquisition Nyquist frequency of 0.25 Hz because neuronal signal is believed to exist above traditional low pass cutoffs (Chen & Glover, 2015; Smith et al., 2013). Subsequently, a condition-specific filter of 0.008 – 0.12Hz was applied to avoid potential artifacts caused by cardiorespiratory noise (Biswal et al., 1996; Biswal et al., 1997). Linear detrending was also performed, and despiking was applied as a final step to remove remaining artifacts in the data (Patel et al., 2014).

Following denoising, average timeseries were extracted from all analyzed ROI seeds using normalized, unsmoothed data. Mean SRCC-corrected, unsmoothed claustrum timeseries were imported as first level covariates and treated as seeds. Seed-based connectivity maps were estimated using the mean timeseries of each unsmoothed ROI seed and all brain voxels from smoothed functional images. Functional connectivity strength was represented by Fisher-transformed bivariate correlation coefficients from a weighted GLM (Nieto-Castanon, 2020), defined separately for each pair of seed and target areas, modeling the association between their BOLD signal timeseries.

Group-level analyses were performed using a GLM (Nieto-Castanon, 2020). For each voxel a separate GLM was estimated, with first-level connectivity measures at this voxel as dependent variables, and groups as independent variables. Voxel-level hypotheses were evaluated using multivariate parametric statistics with random-effects across subjects and sample covariance estimation across multiple measurements. Inferences were performed at the level of individual clusters (groups of contiguous voxels). Cluster-level inferences were based on parametric statistics from Gaussian Random Field theory (Nieto-Castanon, 2020; Worsley et al., 1996). Results were thresholded using a combination of a cluster-forming p < 0.001 voxel-level threshold, and a familywise corrected p-FDR < 0.05 cluster-size threshold (Chumbley et al., 2010).

### Resting State (f)ALFF

Amplitude of Low Frequency Fluctuations (ALFF; Zang et al., 2007) and Fractional Amplitude of Low Frequency Fluctuations (fALFF; Zou et al., 2008) are putative measures of spontaneous neuronal activity. An assessment of the measures’ relative merits determined that ALFF is more susceptible to physiological noise than fALFF near brain cisterns and large blood vessels, but that ALFF is more reliable than fALFF in gray matter regions, making it preferable for group comparisons. It is therefore recommended to report both values (Zou et al., 2010). We opted to initially analyze ALFF and report both measures if an ALFF group difference was detected. We do not believe concerns of physiological noise in ALFF near brain cisterns apply to claustrum analyses. ALFF and fALFF results for non-claustrum ROIs were extracted by masking non-SRCC voxel-wise analyses of unsmoothed resting state images in CONN. RCL and LCL ALFF and fALFF results were extracted by similarly masking voxel-wise analyses in CONN after the respective right or left insular cortex and putamen flanking ROIs were included as additional denoising confounds.

### DWI Analysis in HCP Healthy Individuals

174 participants (106 female, mean age: 29.6) with 7T diffusion data (DWI) were used from the Washington University and University of Minnesota (WU-Minn) S1200 Release from the Human Connectome Project (HCP). Information on inclusion and exclusion criteria are provided elsewhere (Van Essen et al., 2012), but briefly all subjects were healthy with no history of psychological, neurological, or cardiovascular disorders. All 184 subjects with 7T scans within the HCP S1200 release were screened, and 10 were excluded due to missing or corrupted data.

All participant scans were acquired using a Siemens Magnetom scanner with a Nova 32-channel Siemens head coil. DWI scans used EPI obtained over 4 runs with TR 7000ms, TE 71.2ms, slice thickness 1.05mm, and FOV 210mm. Each scan was acquired from two sets of gradient tables, each with a different b-value. Each set contained 65 diffusion-weighted directions, and 6 non-diffusion weighting images (b=0s/mm2; B0) distributed throughout the runs. Diffusion weighting consisted of two shells (b = 1000 and 2000 s/mm2), with equal numbers of acquisitions of each shell throughout each run (Van Essen et al., 2012). Further details about the acquisition can be found at the HCP S1200 Release Reference Manual (https://www.humanconnectome.org/storage/app/media/documentation/s1200/HCP_S1200_Release_Reference_Manual.pdf).

Preprocessed DWI data were downloaded from the HCP (Glasser et al., 2013; Jenkinson et al., 2002), which underwent the generic HCP MR diffusion preprocessing pipeline (v3.19.0), which briefly included: intensity normalization and distortion correction using FSL’s TOPUP and EDDY (v.5.0.10) (Andersson et al., 2003; Andersson & Sotiropoulos, 2015). Scanner gradient nonlinearities were corrected by spatial warping using scanner specific information (Jovicich et al., 2006). Beyond the HCP preprocessing pipeline, all DWI scans underwent tract estimation using FSL’s Bayesian Estimation of Diffusion Parameters Obtained using Sampling Techniques-Crossing Fibers (BEDPOSTX; Behrens et al., 2003; Behrens et al., 2007). BEDPOSTX was run to estimate fiber orientation of each voxel, using a two-fiber model.

Probabilistic tractography was performed using FSL’s Probtrackx2 (Behrens et al., 2003; Behrens et al., 2007) to assess structural connectivity differences between RCL-latDLPFC and RCL-medDLPFC. Each ROI was in MNI152 space and was non-linearly transformed to each individual’s diffusion space using FMRIB’s Non-linear Image Registration Tool (FNIRT; Jenkinson et al., 2012). For each circuit, two tractograms were computed, one from RCL to the target and one from the target to RCL to control for directional biases in acquisition (Van Essen et al., 2013) and fiber fanning (Jeurissen et al., 2019). The modified Euler algorithm was used to generate tractograms with 10,000 streamlines per voxel. To exclude spurious connections, we included exclusion masks (Supplemental Fig. S4) to limit and guide tractography:

i. an insula mask (Cormie et al., 2023),
ii. a thalamus mask, from the Harvard-Oxford subcortical atlas (Desikan et al., 2006; Frazier et al., 2005; Goldstein et al., 2007; Makris et al., 2006),
iii. a mid-sagittal exclusion mask, to remove tracts crossing the midline, and
iv. either a latDLPFC or medDLPFC mask, depending on the circuit studied (i.e., when assessing RCL-latDLPFC connections, a medDLPFC exclusion mask was used).

To control for the direction of streamline propagation, we configured each tractography analysis to propagate toward and not past the DLPFC regions for tracts originating from RCL, or RCL as a terminal point for tracts originating from the DLPFC regions.

Structural connectivity strength is operationalized as the number of streamlines from the seed to the target. As we set Probtrackx2 to send 10,000 streamlines from each ROI seed voxel, the total number of streamlines is 10,000 multiplied by the number of voxels in the seed mask. To correct for bias from different seed volumes, the number of streamlines were divided by 10,000 multiplied by 0.0002 (to avoid spurious connections) multiplied by the number of voxels in the seed of origin. Next, to control for directionality within the same circuit (e.g., RCL-to-latDLPFC and latDLPFC-to-RCL), corrected streamline counts were averaged. Connectivity strength has arbitrary units (a.u.). Group tractograms for both circuits were created by averaging each direction of the circuit for each individual, then adding all participants’ images.

### Dynamic Causal Modeling

DCM was performed in SPM12. Separate DCM analyses were performed for Dataset 1 pain (n = 34 healthy controls, n = 105 patients) and Dataset 1 cognitive task (n = 35 healthy controls, n = 112 patients) scans. All first level models were fully connected, comprising bidirectional RCL←→latDLPFC connectivity and inhibitory self-connections in both regions. Because RCL is upstream of latDLPFC in our hypothesized circuit model, all experimental stimuli (i.e., pain onset, difficult cognitive task onset) were modeled to affect RCL. For each analysis, all estimated first level models were compiled into a fully connected Parametric Empirical Bayes (PEB) model. Because group comparisons were desired, the second level design matrices contained two columns coded to specify 1) the average connectivity parameters among healthy controls and 2) the effect of chronic pain on healthy control parameters (i.e., group differences). The fully connected PEB and all possible reduced models were then searched. Only parameters exceeding the threshold for “positive” evidence (greater than 0.75 posterior probability) were reported.

### Statistical Analyses

FIR results were analyzed via linear mixed effects models in R Statistical Software (v4.2.3; R Core Team 2023) to detect significant signal change compared to average pre-stimulus baseline signal as an effect of timepoint (*y*_*ij*_ [BOLD Signal] = *β*_0_ + *timepoint*_*ij*_ ∗ *β*_1_ + *α*_0*i*_ + *ɛ*_*ij*_), as well as to assess condition effects (*y*_*ij*_ [BOLD Signal] = *β*_0_ + *condition*_*ij*_ ∗ *β*_1_ + *α*_0*i*_ + *ɛ*_*ij*_). Due to the a priori hypothesis of transient claustrum activation at stimulus onset, to minimize comparisons and eliminate selection bias, three “onset” timepoints (t1, t2, t3; treating stimulus onset as t0) and one “block” timepoint (t6 – stimulation block midpoint) were modeled. “Onset” timepoints began after stimulus onset to account for hemodynamic delay in BOLD signal.

Statistical analyses of GLM results were performed in GraphPad Prism 9.4.0.

For DWI analyses, to test whether connectivity strengths were normally distributed, the Kolmogorov-Smirnov test, which is robust for samples >50 participants (Mishra et al., 2019), was used. It was determined that both circuits possessed non-normal distributions (p < 0.05). To compare connectivity in each circuit to a hypothetical median of 0, one-sample Wilcoxon signed-rank tests were performed. To compare the connectivity strength between the two circuits, a non-parametric related-samples Wilcoxon signed-rank test was performed with significance set at p < 0.05.

Multiple comparisons were accounted for in resting state functional connectivity analyses in CONN as mentioned above. Sidak’s multiple comparisons test was used to adjust p-values in ANOVA post-hoc comparisons in GraphPad Prism. When performing multiple statistical tests of other kinds (e.g., one-sample or unpaired t-tests), multiple comparisons correction was performed using Benjamini-Hochberg FDR correction in MATLAB.

### Data and Code Availability

Data and corresponding code are available through request.

---

**Supplemental Figure S1.**
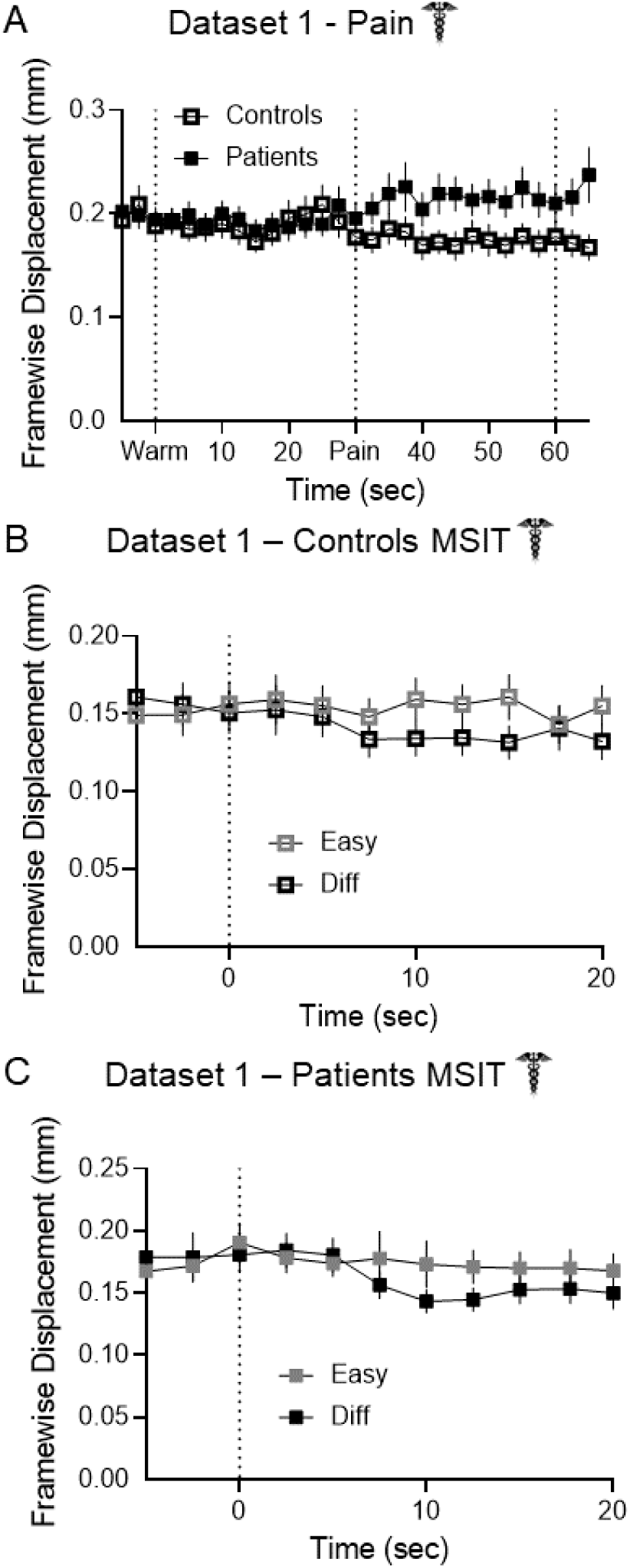
Claustrum signal changes are not attributable to subject movement. **(A)** Neither controls nor patients exhibited transient increases in movement at pain onset. Patients exhibited higher movement values during the pain block than controls, but only controls exhibited statistically significant claustrum signal increases during the pain block. Neither **(B)** controls nor **(C)** patients exhibited transient increases in movement at cognitive task onset, when claustrum activation was detected. No statistical tests were performed on (A) – (C).

**Supplemental Figure S2.**
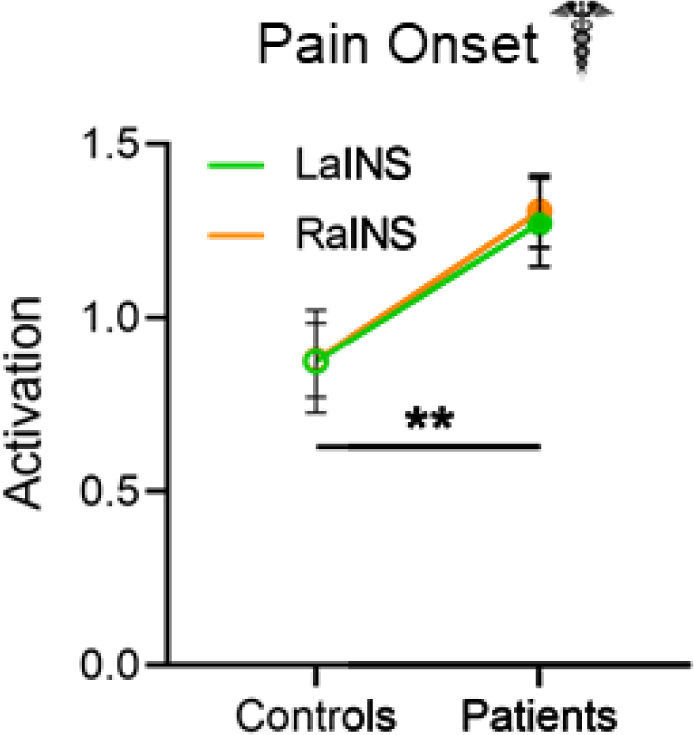
Patients exhibit increased bilateral anterior insula activity at pain onset compared to controls. Two-way ANOVA of bilateral anterior insula activation between groups detected greater activity in patients than controls (main effect of group: F (1, 274) = 7.269, p = 0.007). No significant effect of hemisphere (F (1, 274) = 0.014, p = 0.907) or interaction (F (1, 274) = 0.009, p = 0.924) was detected.

**Supplemental Figure S3.**
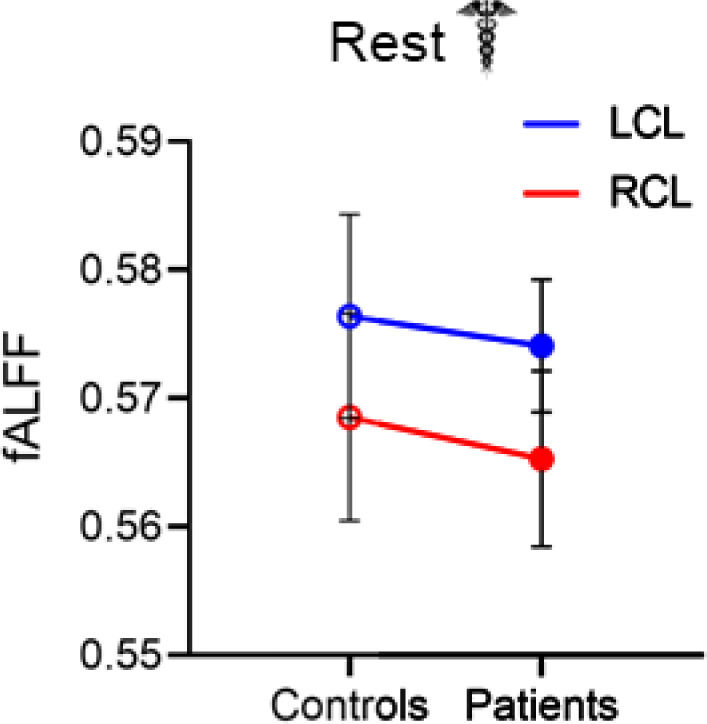
No group or hemisphere differences are present when measuring spontaneous claustrum activity at rest with fALFF. Two-way ANOVA of resting state fALFF between controls and migraine patients detected no main effect of group (F (1, 280) = 0.109, p = P=0.742), hemisphere (F (1, 280) = 0.991, p = P=0.320), or their interaction (F (1, 280) = 0.003, p = P=0.956).

**Supplemental Figure S4.**
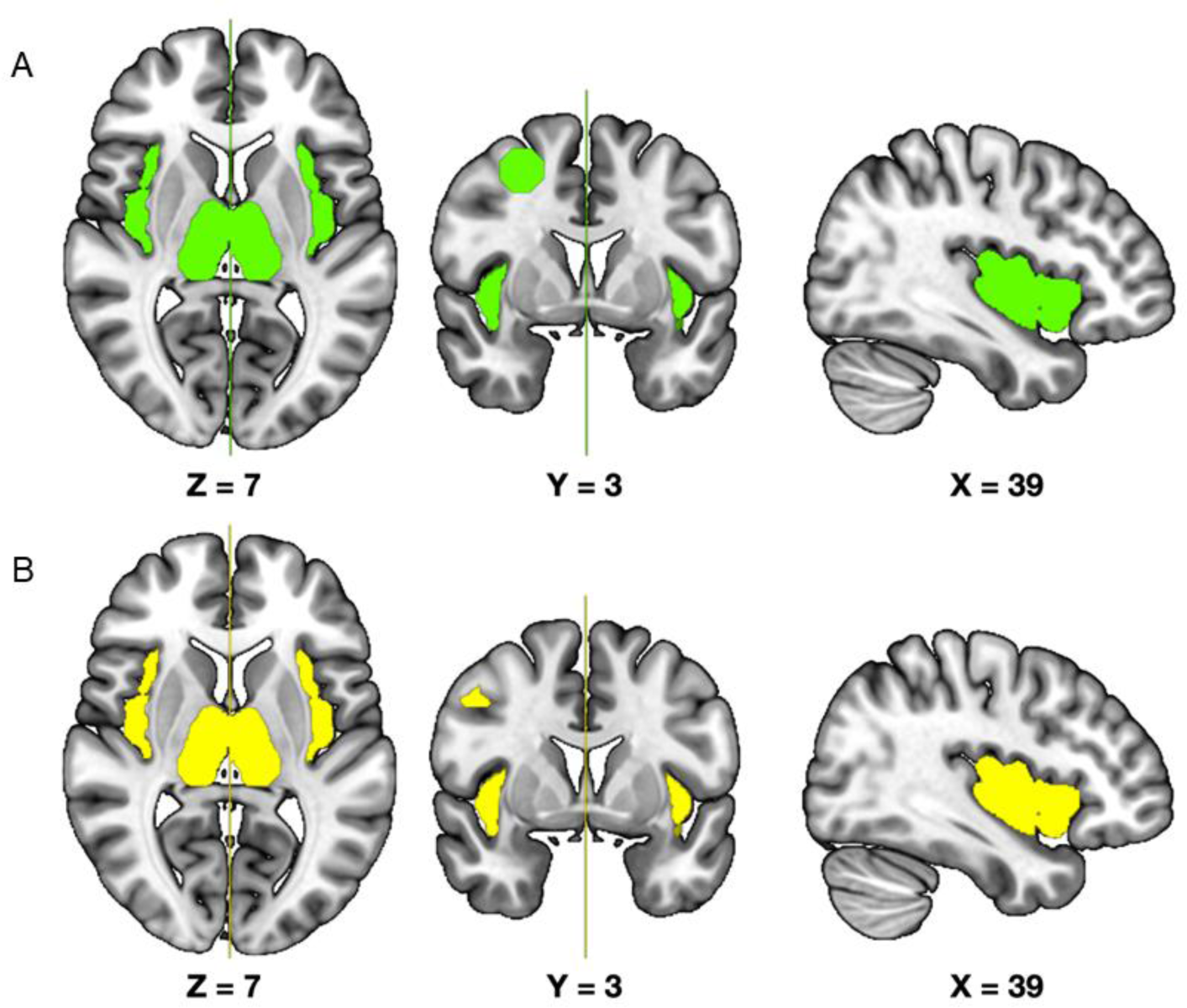
DWI analysis exclusion masks. Exclusion masks used in DWI analyses between **(A)** RCL and latDLPFC (excluding medDLPFC), and **(B)** RCL and medDLPFC (excluding latDLPFC).

**Supplemental Table 1.** Clusters exhibiting significant differences between patients and controls in cognitive task-associated activity (PLS). All clusters display greater activity in patients than controls.

| Cluster # | Peak Location | Bootstrap Ratio | Approx. P-Value | Cluster Size (voxels) |
| --- | --- | --- | --- | --- |
| 1 | 52, 0, 44 | 6.4332 | 0.0000 | 261 |
| 2 | -36, -78, -14 | 4.8469 | 0.0000 | 110 |
| 3 | 32, -64, -12 | 4.7768 | 0.0000 | 231 |
| 4 | -28, 26, -10 | 4.7223 | 0.0000 | 67 |
| 5 | 8, -8, 20 | 4.6402 | 0.0000 | 158 |
| 6 | 30, 22, -8 | 4.4100 | 0.0000 | 110 |
| 7 | -6, -72, -36 | 4.3793 | 0.0000 | 85 |
| 8 | 30, -84, -6 | 4.3370 | 0.0000 | 52 |
| 9 | 32, -46, -34 | 4.1539 | 0.0000 | 50 |
| 10 | 12, -42, -30 | 4.0413 | 0.0001 | 52 |
| 11 | 4, -60, -36 | 3.9191 | 0.0001 | 55 |
| 12 | -6, 38, 22 | 3.9106 | 0.0001 | 73 |

## Notes

### Competing Interest Statement

The authors have declared no competing interest.

## References

Abdallah M, Zanitti GE, Iovene V, Wassermann D. Functional gradients in the human lateral prefrontal cortex revealed by a comprehensive coordinate-based meta-analysis. Elife. 2022 Sep 28;11:e76926. doi: 10.7554/eLife.76926. PMID: 36169404; PMCID: PMC9578708.

Allen M, Dietz M, Blair KS, van Beek M, Rees G, Vestergaard-Poulsen P, Lutz A, Roepstorff A. Cognitive-affective neural plasticity following active-controlled mindfulness intervention. J Neurosci. 2012 Oct 31;32(44):15601–10. doi: 10.1523/JNEUROSCI.2957-12.2012. PMID: 23115195; PMCID: PMC4569704.

Andersson JL, Skare S, Ashburner J. How to correct susceptibility distortions in spin-echo echo-planar images: application to diffusion tensor imaging. Neuroimage. 2003 Oct;20(2):870–88. doi: 10.1016/S1053-8119(03)00336-7. PMID: 14568458.

Andersson JL, Sotiropoulos SN. Non-parametric representation and prediction of single- and multi-shell diffusion-weighted MRI data using Gaussian processes. Neuroimage. 2015 Nov 15;122:166–76. doi: 10.1016/j.neuroimage.2015.07.067. Epub 2015 Jul 30. PMID: 26236030; PMCID: PMC4627362.

Atlas LY, Bolger N, Lindquist MA, Wager TD. Brain mediators of predictive cue effects on perceived pain. J Neurosci. 2010 Sep 29;30(39):12964–77. doi: 10.1523/JNEUROSCI.0057-10.2010. PMID: 20881115; PMCID: PMC2966558.

Atlan G, Terem A, Peretz-Rivlin N, Sehrawat K, Gonzales BJ, Pozner G, Tasaka GI, Goll Y, Refaeli R, Zviran O, Lim BK, Groysman M, Goshen I, Mizrahi A, Nelken I, Citri A. The Claustrum Supports Resilience to Distraction. Curr Biol. 2018 Sep 10;28(17):2752–2762.e7. doi: 10.1016/j.cub.2018.06.068. Epub 2018 Aug 16. PMID: 30122531; PMCID: PMC6485402.

Baker KS, Gibson S, Georgiou-Karistianis N, Roth RM, Giummarra MJ. Everyday Executive Functioning in Chronic Pain: Specific Deficits in Working Memory and Emotion Control, Predicted by Mood, Medications, and Pain Interference. Clin J Pain. 2016 Aug;32(8):673–80. doi: 10.1097/AJP.0000000000000313. PMID: 26626294.

Baliki MN, Geha PY, Apkarian AV, Chialvo DR. Beyond feeling: chronic pain hurts the brain, disrupting the default-mode network dynamics. J Neurosci. 2008 Feb 6;28(6):1398–403. doi: 10.1523/JNEUROSCI.4123-07.2008. PMID: 18256259; PMCID: PMC6671589.

Barbey AK, Koenigs M, Grafman J. Dorsolateral prefrontal contributions to human working memory. Cortex. 2013 May;49(5):1195–205. doi: 10.1016/j.cortex.2012.05.022. Epub 2012 Jun 16. PMID: 22789779; PMCID: PMC3495093.

Barrett FS, Krimmel SR, Griffiths RR, Seminowicz DA, Mathur BN. Psilocybin acutely alters the functional connectivity of the claustrum with brain networks that support perception, memory, and attention. Neuroimage. 2020 Sep;218:116980. doi: 10.1016/j.neuroimage.2020.116980. Epub 2020 May 23. PMID: 32454209.

Bassett DS, Bullmore E, Verchinski BA, Mattay VS, Weinberger DR, Meyer-Lindenberg A. Hierarchical organization of human cortical networks in health and schizophrenia. J Neurosci. 2008 Sep 10;28(37):9239–48. doi: 10.1523/JNEUROSCI.1929-08.2008. PMID: 18784304; PMCID: PMC2878961.

Behrens TE, Berg HJ, Jbabdi S, Rushworth MF, Woolrich MW. Probabilistic diffusion tractography with multiple fibre orientations: What can we gain? Neuroimage. 2007 Jan 1;34(1):144–55. doi: 10.1016/j.neuroimage.2006.09.018. Epub 2006 Oct 27. PMID: 17070705; PMCID: PMC7116582.

Behrens TE, Woolrich MW, Jenkinson M, Johansen-Berg H, Nunes RG, Clare S, Matthews PM, Brady JM, Smith SM. Characterization and propagation of uncertainty in diffusion-weighted MR imaging. Magn Reson Med. 2003 Nov;50(5):1077–88. doi: 10.1002/mrm.10609. PMID: 14587019.

Behzadi Y, Restom K, Liau J, Liu TT. A component based noise correction method (CompCor) for BOLD and perfusion based fMRI. Neuroimage. 2007 Aug 1;37(1):90–101. doi: 10.1016/j.neuroimage.2007.04.042. Epub 2007 May 3. PMID: 17560126; PMCID: PMC2214855.

Berryman C, Stanton TR, Bowering KJ, Tabor A, McFarlane A, Moseley GL. Do people with chronic pain have impaired executive function? A meta-analytical review. Clin Psychol Rev. 2014 Nov;34(7):563–79. doi: 10.1016/j.cpr.2014.08.003. Epub 2014 Sep 16. PMID: 25265056.

Bidet-Caulet A, Buchanan KG, Viswanath H, Black J, Scabini D, Bonnet-Brilhault F, Knight RT. Impaired Facilitatory Mechanisms of Auditory Attention After Damage of the Lateral Prefrontal Cortex. Cereb Cortex. 2015 Nov;25(11):4126–34. doi: 10.1093/cercor/bhu131. Epub 2014 Jun 12. PMID: 24925773; PMCID: PMC4626830.

Biswal B, DeYoe EA, Hyde JS. Reduction of physiological fluctuations in fMRI using digital filters. Magn Reson Med. 1996 Jan;35(1):107–13. doi: 10.1002/mrm.1910350114. PMID: 8771028.

Biswal B, Hudetz AG, Yetkin FZ, Haughton VM, Hyde JS. Hypercapnia reversibly suppresses low-frequency fluctuations in the human motor cortex during rest using echo-planar MRI. J Cereb Blood Flow Metab. 1997 Mar;17(3):301–8. doi: 10.1097/00004647-199703000-00007. PMID: 9119903.

Buhle JT, Silvers JA, Wager TD, Lopez R, Onyemekwu C, Kober H, Weber J, Ochsner KN. Cognitive reappraisal of emotion: a meta-analysis of human neuroimaging studies. Cereb Cortex. 2014 Nov;24(11):2981–90. doi: 10.1093/cercor/bht154. Epub 2013 Jun 13. PMID: 23765157; PMCID: PMC4193464.

Buhle J, Wager TD. Performance-dependent inhibition of pain by an executive working memory task. Pain. 2010 Apr;149(1):19–26. doi: 10.1016/j.pain.2009.10.027. Epub 2010 Feb 2. PMID: 20129735; PMCID: PMC4229048.

Bush G, Shin LM, Holmes J, Rosen BR, Vogt BA. The Multi-Source Interference Task: validation study with fMRI in individual subjects. Mol Psychiatry. 2003 Jan;8(1):60–70. doi: 10.1038/sj.mp.4001217. PMID: 12556909.

Carhart-Harris RL, Friston KJ. REBUS and the Anarchic Brain: Toward a Unified Model of the Brain Action of Psychedelics. Pharmacol Rev. 2019 Jul;71(3):316–344. doi: 10.1124/pr.118.017160. PMID: 31221820; PMCID: PMC6588209.

Castellanos JP, Woolley C, Bruno KA, Zeidan F, Halberstadt A, Furnish T. Chronic pain and psychedelics: a review and proposed mechanism of action. Reg Anesth Pain Med. 2020 Jul;45(7):486–494. doi: 10.1136/rapm-2020-101273. Epub 2020 May 4. PMID: 32371500.

Čeko M, Gracely JL, Fitzcharles MA, Seminowicz DA, Schweinhardt P, Bushnell MC. Is a Responsive Default Mode Network Required for Successful Working Memory Task Performance? J Neurosci. 2015 Aug 19;35(33):11595–605. doi: 10.1523/JNEUROSCI.0264-15.2015. PMID: 26290236; PMCID: PMC4540797.

Chen JE, Glover GH. BOLD fractional contribution to resting-state functional connectivity above 0.1 Hz. Neuroimage. 2015 Feb 15;107:207–218. doi: 10.1016/j.neuroimage.2014.12.012. Epub 2014 Dec 12. PMID: 25497686; PMCID: PMC4318656.

Chumbley J, Worsley K, Flandin G, Friston K. Topological FDR for neuroimaging. Neuroimage. 2010 Feb 15;49(4):3057–64. doi: 10.1016/j.neuroimage.2009.10.090. Epub 2009 Nov 24. PMID: 19944173; PMCID: PMC3221040.

Cormie MA, Kaya B, Hadjis GE, Mouseli P, Moayedi M. Insula-cingulate structural and functional connectivity: an ultra-high field MRI study. Cereb Cortex. 2023 Aug 23;33(17):9787–9801. doi: 10.1093/cercor/bhad244. PMID: 37429832.

Desikan RS, Ségonne F, Fischl B, Quinn BT, Dickerson BC, Blacker D, Buckner RL, Dale AM, Maguire RP, Hyman BT, Albert MS, Killiany RJ. An automated labeling system for subdividing the human cerebral cortex on MRI scans into gyral based regions of interest. Neuroimage. 2006 Jul 1;31(3):968–80. doi: 10.1016/j.neuroimage.2006.01.021. Epub 2006 Mar 10. PMID: 16530430.

Dosenbach NU, Fair DA, Cohen AL, Schlaggar BL, Petersen SE. A dual-networks architecture of top-down control. Trends Cogn Sci. 2008 Mar;12(3):99–105. doi: 10.1016/j.tics.2008.01.001. Epub 2008 Feb 11. PMID: 18262825; PMCID: PMC3632449.

Dosenbach NU, Fair DA, Miezin FM, Cohen AL, Wenger KK, Dosenbach RA, Fox MD, Snyder AZ, Vincent JL, Raichle ME, Schlaggar BL, Petersen SE. Distinct brain networks for adaptive and stable task control in humans. Proc Natl Acad Sci U S A. 2007 Jun 26;104(26):11073–8. doi: 10.1073/pnas.0704320104. Epub 2007 Jun 18. PMID: 17576922; PMCID: PMC1904171.

Doss MK, Madden MB, Gaddis A, Nebel MB, Griffiths RR, Mathur BN, Barrett FS. Models of psychedelic drug action: modulation of cortical-subcortical circuits. Brain. 2022 Apr 18;145(2):441–456. doi: 10.1093/brain/awab406. PMID: 34897383; PMCID: PMC9014750.

Eccleston C, Crombez G. Pain demands attention: a cognitive-affective model of the interruptive function of pain. Psychol Bull. 1999 May;125(3):356–66. doi: 10.1037/0033-2909.125.3.356. PMID: 10349356.

Eck J, Richter M, Straube T, Miltner WHR, Weiss T. Affective brain regions are activated during the processing of pain-related words in migraine patients. Pain. 2011 May;152(5):1104–1113. doi: 10.1016/j.pain.2011.01.026. Epub 2011 Mar 5. PMID: 21377797.

Frazier JA, Chiu S, Breeze JL, Makris N, Lange N, Kennedy DN, Herbert MR, Bent EK, Koneru VK, Dieterich ME, Hodge SM, Rauch SL, Grant PE, Cohen BM, Seidman LJ, Caviness VS, Biederman J. Structural brain magnetic resonance imaging of limbic and thalamic volumes in pediatric bipolar disorder. Am J Psychiatry. 2005 Jul;162(7):1256–65. doi: 10.1176/appi.ajp.162.7.1256. PMID: 15994707.

Friston KJ, Harrison L, Penny W. Dynamic causal modelling. Neuroimage. 2003 Aug;19(4):1273-302. doi: 10.1016/s1053-8119(03)00202-7. PMID: 12948688.

Glasser MF, Sotiropoulos SN, Wilson JA, Coalson TS, Fischl B, Andersson JL, Xu J, Jbabdi S, Webster M, Polimeni JR, Van Essen DC, Jenkinson M; WU-Minn HCP Consortium. The minimal preprocessing pipelines for the Human Connectome Project. Neuroimage. 2013 Oct 15;80:105–24. doi: 10.1016/j.neuroimage.2013.04.127. Epub 2013 May 11. PMID: 23668970; PMCID: PMC3720813.

Goldstein JM, Seidman LJ, Makris N, Ahern T, O’Brien LM, Caviness VS Jr, Kennedy DN, Faraone SV, Tsuang MT. Hypothalamic abnormalities in schizophrenia: sex effects and genetic vulnerability. Biol Psychiatry. 2007 Apr 15;61(8):935–45. doi: 10.1016/j.biopsych.2006.06.027. Epub 2006 Oct 13. PMID: 17046727.

Gorgolewski KJ, Mendes N, Wilfling D, Wladimirow E, Gauthier CJ, Bonnen T, Ruby FJ, Trampel R, Bazin PL, Cozatl R, Smallwood J, Margulies DS. A high resolution 7-Tesla resting-state fMRI test-retest dataset with cognitive and physiological measures. Sci Data. 2015 Jan 20;2:140054. doi: 10.1038/sdata.2014.54. PMID: 25977805; PMCID: PMC4412153.

Gracely RH, Geisser ME, Giesecke T, Grant MA, Petzke F, Williams DA, Clauw DJ. Pain catastrophizing and neural responses to pain among persons with fibromyalgia. Brain. 2004 Apr;127(Pt 4):835–43. doi: 10.1093/brain/awh098. Epub 2004 Feb 11. PMID: 14960499.

Graff-Guerrero A, González-Olvera J, Fresán A, Gómez-Martín D, Méndez-Núñez JC, Pellicer F. Repetitive transcranial magnetic stimulation of dorsolateral prefrontal cortex increases tolerance to human experimental pain. Brain Res Cogn Brain Res. 2005 Sep;25(1):153–60. doi: 10.1016/j.cogbrainres.2005.05.002. PMID: 15935625.

Greicius MD, Krasnow B, Reiss AL, Menon V. Functional connectivity in the resting brain: a network analysis of the default mode hypothesis. Proc Natl Acad Sci U S A. 2003 Jan 7;100(1):253–8. doi: 10.1073/pnas.0135058100. Epub 2002 Dec 27. PMID: 12506194; PMCID: PMC140943.

Gu L, Shu H, Wang Y. Functional brain alterations in migraine patients: an activation likelihood estimation study. Neurol Res. 2023 Apr 5:1–8. doi: 10.1080/01616412.2023.2199377. Epub ahead of print. PMID: 37019685.

Hallquist MN, Hwang K, Luna B. The nuisance of nuisance regression: spectral misspecification in a common approach to resting-state fMRI preprocessing reintroduces noise and obscures functional connectivity. Neuroimage. 2013 Nov 15;82:208–25. doi: 10.1016/j.neuroimage.2013.05.116. Epub 2013 Jun 6. PMID: 23747457; PMCID: PMC3759585.

Hashmi JA, Baliki MN, Huang L, Baria AT, Torbey S, Hermann KM, Schnitzer TJ, Apkarian AV. Shape shifting pain: chronification of back pain shifts brain representation from nociceptive to emotional circuits. Brain. 2013 Sep;136(Pt 9):2751–68. doi: 10.1093/brain/awt211. PMID: 23983029; PMCID: PMC3754458.

Hubbard CS, Khan SA, Keaser ML, Mathur VA, Goyal M, Seminowicz DA. Altered Brain Structure and Function Correlate with Disease Severity and Pain Catastrophizing in Migraine Patients. eNeuro. 2014 Dec 30;1(1):e20.14. doi: 10.1523/ENEURO.0006-14.2014. PMID: 25893216; PMCID: PMC4399775.

Jenkinson M, Bannister P, Brady M, Smith S. Improved optimization for the robust and accurate linear registration and motion correction of brain images. Neuroimage. 2002 Oct;17(2):825–41. doi: 10.1016/s1053-8119(02)91132-8. PMID: 12377157.

Jenkinson M, Beckmann CF, Behrens TE, Woolrich MW, Smith SM. FSL. Neuroimage. 2012 Aug 15;62(2):782-90. doi: 10.1016/j.neuroimage.2011.09.015. Epub 2011 Sep 16. PMID: 21979382.

Jeurissen B, Descoteaux M, Mori S, Leemans A. Diffusion MRI fiber tractography of the brain. NMR Biomed. 2019 Apr;32(4):e3785. doi: 10.1002/nbm.3785. Epub 2017 Sep 25. PMID: 28945294.

Jovicich J, Czanner S, Greve D, Haley E, van der Kouwe A, Gollub R, Kennedy D, Schmitt F, Brown G, Macfall J, Fischl B, Dale A. Reliability in multi-site structural MRI studies: effects of gradient non-linearity correction on phantom and human data. Neuroimage. 2006 Apr 1;30(2):436–43. doi: 10.1016/j.neuroimage.2005.09.046. Epub 2005 Nov 21. PMID: 16300968.

Jung J, Lambon Ralph MA, Jackson RL. Subregions of DLPFC Display Graded yet Distinct Structural and Functional Connectivity. J Neurosci. 2022 Apr 13;42(15):3241–3252. doi: 10.1523/JNEUROSCI.1216-21.2022. Epub 2022 Mar 1. PMID: 35232759; PMCID: PMC8994544.

Koechlin E, Ody C, Kouneiher F. The architecture of cognitive control in the human prefrontal cortex. Science. 2003 Nov 14;302(5648):1181-5. doi: 10.1126/science.1088545. PMID: 14615530.

Krimmel SR, White MG, Panicker MH, Barrett FS, Mathur BN, Seminowicz DA. Resting state functional connectivity and cognitive task-related activation of the human claustrum. Neuroimage. 2019 Aug 1;196:59–67. doi: 10.1016/j.neuroimage.2019.03.075. Epub 2019 Apr 4. PMID: 30954711; PMCID: PMC6629463.

Krummenacher P, Candia V, Folkers G, Schedlowski M, Schönbächler G. Prefrontal cortex modulates placebo analgesia. Pain. 2010 Mar;148(3):368–374. doi: 10.1016/j.pain.2009.09.033. Epub 2009 Oct 28. PMID: 19875233.

Landrø NI, Fors EA, Våpenstad LL, Holthe Ø, Stiles TC, Borchgrevink PC. The extent of neurocognitive dysfunction in a multidisciplinary pain centre population. Is there a relation between reported and tested neuropsychological functioning? Pain. 2013 Jul;154(7):972–7. doi: 10.1016/j.pain.2013.01.013. Epub 2013 Feb 8. PMID: 23473784.

Lesh TA, Niendam TA, Minzenberg MJ, Carter CS. Cognitive control deficits in schizophrenia: mechanisms and meaning. Neuropsychopharmacology. 2011 Jan;36(1):316–38. doi: 10.1038/npp.2010.156. Epub 2010 Sep 15. PMID: 20844478; PMCID: PMC3052853.

Lorenz J, Minoshima S, Casey KL. Keeping pain out of mind: the role of the dorsolateral prefrontal cortex in pain modulation. Brain. 2003 May;126(Pt 5):1079–91. doi: 10.1093/brain/awg102. PMID: 12690048.

McIntosh AR, Bookstein FL, Haxby JV, Grady CL. Spatial pattern analysis of functional brain images using partial least squares. Neuroimage. 1996 Jun;3(3 Pt 1):143-57. doi: 10.1006/nimg.1996.0016. PMID: 9345485.

Madden MB, Stewart BW, White MG, Krimmel SR, Qadir H, Barrett FS, Seminowicz DA, Mathur BN. A role for the claustrum in cognitive control. Trends Cogn Sci. 2022 Dec;26(12):1133–1152. doi: 10.1016/j.tics.2022.09.006. Epub 2022 Sep 30. PMID: 36192309; PMCID: PMC9669149.

Makris N, Goldstein JM, Kennedy D, Hodge SM, Caviness VS, Faraone SV, Tsuang MT, Seidman LJ. Decreased volume of left and total anterior insular lobule in schizophrenia. Schizophr Res. 2006 Apr;83(2-3):155–71. doi: 10.1016/j.schres.2005.11.020. Epub 2006 Jan 31. PMID: 16448806.

Markowitsch HJ, Irle E, Bang-Olsen R, Flindt-Egebak P. Claustral efferents to the cat’s limbic cortex studied with retrograde and anterograde tracing techniques. Neuroscience. 1984 Jun;12(2):409–25. doi: 10.1016/0306-4522(84)90062-9. PMID: 6462456.

Mathur VA, Khan SA, Keaser ML, Hubbard CS, Goyal M, Seminowicz DA. Altered cognition-related brain activity and interactions with acute pain in migraine. Neuroimage Clin. 2015 Jan 10;7:347–58. doi: 10.1016/j.nicl.2015.01.003. PMID: 25610798; PMCID: PMC4297882.

Brett M, Anton JL, Valabregue R, Poline JB. Region of interest analysis using an SPM toolbox [abstract] Presented at the 8th International Conference on Functional Mapping of the Human Brain, June 2–6, 2002, Sendai, Japan. Available on CD-ROM in NeuroImage, Vol 16, No 2.

Melzack, R., Casey, K (1968). Sensory, motivational, and central control determinants of pain: A new conceptual model. In The Skin Senses, D.R. Kenshalo, ed. (Springfield, IL: Thomas) pp. 423– 443.

Menon V, D’Esposito M. The role of PFC networks in cognitive control and executive function. Neuropsychopharmacology. 2022 Jan;47(1):90–103. doi: 10.1038/s41386-021-01152-w. Epub 2021 Aug 18. PMID: 34408276; PMCID: PMC8616903.

Mishra P, Pandey CM, Singh U, Gupta A, Sahu C, Keshri A. Descriptive statistics and normality tests for statistical data. Ann Card Anaesth. 2019 Jan-Mar;22(1):67-72. doi: 10.4103/aca.ACA_157_18. PMID: 30648682; PMCID: PMC6350423.

Murphy K, Fox MD. Towards a consensus regarding global signal regression for resting state functional connectivity MRI. Neuroimage. 2017 Jul 1;154:169–173. doi: 10.1016/j.neuroimage.2016.11.052. Epub 2016 Nov 22. PMID: 27888059; PMCID: PMC5489207.

Muschelli J, Nebel MB, Caffo BS, Barber AD, Pekar JJ, Mostofsky SH. Reduction of motion-related artifacts in resting state fMRI using aCompCor. Neuroimage. 2014 Aug 1;96:22–35. doi: 10.1016/j.neuroimage.2014.03.028. Epub 2014 Mar 18. PMID: 24657780; PMCID: PMC4043948.

Nee DE, D’Esposito M. The hierarchical organization of the lateral prefrontal cortex. Elife. 2016 Mar 21;5:e12112. doi: 10.7554/eLife.12112. PMID: 26999822; PMCID: PMC4811776.

Nieto-Castanon, A. (2020). Handbook of functional connectivity Magnetic Resonance Imaging methods in CONN. Boston, MA: Hilbert Press.

Ntamati NR, Acuña MA, Nevian T. Pain-induced adaptations in the claustro-cingulate pathway. Cell Rep. 2023 May 30;42(5):112506. doi: 10.1016/j.celrep.2023.112506. Epub 2023 May 13. PMID: 37182208.

Paul K, Tik M, Hahn A, Sladky R, Geissberger N, Wirth EM, Kranz GS, Pfabigan DM, Kraus C, Lanzenberger R, Lamm C, Windischberger C. Give me a pain that I am used to: distinct habituation patterns to painful and non-painful stimulation. Sci Rep. 2021 Nov 25;11(1):22929. doi: 10.1038/s41598-021-01881-4. PMID: 34824311; PMCID: PMC8617189.

Patel AX, Kundu P, Rubinov M, Jones PS, Vértes PE, Ersche KD, Suckling J, Bullmore ET. A wavelet method for modeling and despiking motion artifacts from resting-state fMRI time series. Neuroimage. 2014 Jul 15;95(100):287–304. doi: 10.1016/j.neuroimage.2014.03.012. Epub 2014 Mar 21. PMID: 24657353; PMCID: PMC4068300.

Petrides M. Lateral prefrontal cortex: architectonic and functional organization. Philos Trans R Soc Lond B Biol Sci. 2005 Apr 29;360(1456):781-95. doi: 10.1098/rstb.2005.1631. PMID: 15937012; PMCID: PMC1569489.

Qadir H, Krimmel SR, Mu C, Poulopoulos A, Seminowicz DA, Mathur BN. Structural Connectivity of the Anterior Cingulate Cortex, Claustrum, and the Anterior Insula of the Mouse. Front Neuroanat. 2018 Nov 26;12:100. doi: 10.3389/fnana.2018.00100. PMID: 30534060; PMCID: PMC6276828.

Qadir H, Stewart BW, VanRyzin JW, Wu Q, Chen S, Seminowicz DA, Mathur BN. The mouse claustrum synaptically connects cortical network motifs. Cell Rep. 2022 Dec 20;41(12):111860. doi: 10.1016/j.celrep.2022.111860. PMID: 36543121; PMCID: PMC9838879.

R Core Team (2023). R: A language and environment for statistical computing. R Foundation for Statistical Computing, Vienna, Austria. URL https://www.R-project.org/.

Rahnev D, Nee DE, Riddle J, Larson AS, D’Esposito M. Causal evidence for frontal cortex organization for perceptual decision making. Proc Natl Acad Sci U S A. 2016 May 24;113(21):6059–64. doi: 10.1073/pnas.1522551113. Epub 2016 May 9. PMID: 27162349; PMCID: PMC4889369.

Reser DH, Richardson KE, Montibeller MO, Zhao S, Chan JM, Soares JG, Chaplin TA, Gattass R, Rosa MG. Claustrum projections to prefrontal cortex in the capuchin monkey (Cebus apella). Front Syst Neurosci. 2014 Jul 3;8:123. doi: 10.3389/fnsys.2014.00123. PMID: 25071475; PMCID: PMC4079979.

Rombouts SA, Barkhof F, Goekoop R, Stam CJ, Scheltens P. Altered resting state networks in mild cognitive impairment and mild Alzheimer’s disease: an fMRI study. Hum Brain Mapp. 2005 Dec;26(4):231–9. doi: 10.1002/hbm.20160. PMID: 15954139; PMCID: PMC6871685.

Schindler EAD, Sewell RA, Gottschalk CH, Luddy C, Flynn LT, Lindsey H, Pittman BP, Cozzi NV, D’Souza DC. Exploratory Controlled Study of the Migraine-Suppressing Effects of Psilocybin. Neurotherapeutics. 2021 Jan;18(1):534–543. doi: 10.1007/s13311-020-00962-y. Epub 2020 Nov 12. PMID: 33184743; PMCID: PMC8116458.

Seeley WW, Menon V, Schatzberg AF, Keller J, Glover GH, Kenna H, Reiss AL, Greicius MD. Dissociable intrinsic connectivity networks for salience processing and executive control. J Neurosci. 2007 Feb 28;27(9):2349–56. doi: 10.1523/JNEUROSCI.5587-06.2007. PMID: 17329432; PMCID: PMC2680293.

Seminowicz DA, Burrowes SAB, Kearson A, Zhang J, Krimmel SR, Samawi L, Furman AJ, Keaser ML, Gould NF, Magyari T, White L, Goloubeva O, Goyal M, Peterlin BL, Haythornthwaite JA. Enhanced mindfulness-based stress reduction in episodic migraine: a randomized clinical trial with magnetic resonance imaging outcomes. Pain. 2020 Aug;161(8):1837–1846. doi: 10.1097/j.pain.0000000000001860. Epub 2020 Mar 13. PMID: 32701843; PMCID: PMC7487005.

Seminowicz DA, Davis KD. Cortical responses to pain in healthy individuals depends on pain catastrophizing. Pain. 2006 Feb;120(3):297–306. doi: 10.1016/j.pain.2005.11.008. Epub 2006 Jan 19. PMID: 16427738.

Seminowicz DA, Davis KD. Pain enhances functional connectivity of a brain network evoked by performance of a cognitive task. J Neurophysiol. 2007 May;97(5):3651–9. doi: 10.1152/jn.01210.2006. Epub 2007 Feb 21. PMID: 17314240.

Seminowicz DA, Wideman TH, Naso L, Hatami-Khoroushahi Z, Fallatah S, Ware MA, Jarzem P, Bushnell MC, Shir Y, Ouellet JA, Stone LS. Effective treatment of chronic low back pain in humans reverses abnormal brain anatomy and function. J Neurosci. 2011 May 18;31(20):7540–50. doi: 10.1523/JNEUROSCI.5280-10.2011. PMID: 21593339; PMCID: PMC6622603.

Shackman AJ, Salomons TV, Slagter HA, Fox AS, Winter JJ, Davidson RJ. The integration of negative affect, pain and cognitive control in the cingulate cortex. Nat Rev Neurosci. 2011 Mar;12(3):154–67. doi: 10.1038/nrn2994. PMID: 21331082; PMCID: PMC3044650.

Smith SM, Beckmann CF, Andersson J, Auerbach EJ, Bijsterbosch J, Douaud G, Duff E, Feinberg DA, Griffanti L, Harms MP, Kelly M, Laumann T, Miller KL, Moeller S, Petersen S, Power J, Salimi-Khorshidi G, Snyder AZ, Vu AT, Woolrich MW, Xu J, Yacoub E, Uğurbil K, Van Essen DC, Glasser MF; WU-Minn HCP Consortium. Resting-state fMRI in the Human Connectome Project. Neuroimage. 2013 Oct 15;80:144–68. doi: 10.1016/j.neuroimage.2013.05.039. Epub 2013 May 20. PMID: 23702415; PMCID: PMC3720828.

Stiernman LJ, Grill F, Hahn A, Rischka L, Lanzenberger R, Panes Lundmark V, Riklund K, Axelsson J, Rieckmann A. Dissociations between glucose metabolism and blood oxygenation in the human default mode network revealed by simultaneous PET-fMRI. Proc Natl Acad Sci U S A. 2021 Jul 6;118(27):e2021913118. doi: 10.1073/pnas.2021913118. PMID: 34193521; PMCID: PMC8271663.

Stiernman L, Grill F, McNulty C, Bahrd P, Panes Lundmark V, Axelsson J, Salami A, Rieckmann A. Widespread fMRI BOLD Signal Overactivations during Cognitive Control in Older Adults Are Not Matched by Corresponding Increases in fPET Glucose Metabolism. J Neurosci. 2023 Apr 5;43(14):2527–2536. doi: 10.1523/JNEUROSCI.1331-22.2023. Epub 2023 Mar 3. PMID: 36868855; PMCID: PMC10082451.

Tanné-Gariépy J, Boussaoud D, Rouiller EM. Projections of the claustrum to the primary motor, premotor, and prefrontal cortices in the macaque monkey. J Comp Neurol. 2002 Dec 9;454(2):140–57. doi: 10.1002/cne.10425. PMID: 12412139.

Taylor KS, Seminowicz DA, Davis KD. Two systems of resting state connectivity between the insula and cingulate cortex. Hum Brain Mapp. 2009 Sep;30(9):2731–45. doi: 10.1002/hbm.20705. PMID: 19072897; PMCID: PMC6871122.

Torgerson CM, Irimia A, Goh SY, Van Horn JD. The DTI connectivity of the human claustrum. Hum Brain Mapp. 2015 Mar;36(3):827–38. doi: 10.1002/hbm.22667. Epub 2014 Oct 23. PMID: 25339630; PMCID: PMC4324054.

Ushida T, Ikemoto T, Tanaka S, Shinozaki J, Taniguchi S, Murata Y, McLaughlin M, Arai YC, Tamura Y. Virtual needle pain stimuli activates cortical representation of emotions in normal volunteers. Neurosci Lett. 2008 Jul 4;439(1):7–12. doi: 10.1016/j.neulet.2008.04.085. Epub 2008 Apr 30. PMID: 18502045.

Van Essen DC, Dierker DL. Surface-based and probabilistic atlases of primate cerebral cortex. Neuron. 2007 Oct 25;56(2):209–25. doi: 10.1016/j.neuron.2007.10.015. PMID: 17964241.

Van Essen DC, Jbabdi S, Sotiropoulos SN, Chen C, Dikranian K, Coalson T, Harwell J, Behrens TEJ, Glasser MF. Mapping Connections in Humans and Non-Human Primates. Aspirations and Challenges for Diffusion Imaging. In Diffusion MRI: From Quantitative Measurement to In vivo Neuroanatomy: Second Edition. 2013. p. 337–358. Elsevier Inc.. 10.1016/B978-0-12-396460-1.00016-0

Van Essen DC, Smith SM, Barch DM, Behrens TE, Yacoub E, Ugurbil K; WU-Minn HCP Consortium. The WU-Minn Human Connectome Project: an overview. Neuroimage. 2013 Oct 15;80:62–79. doi: 10.1016/j.neuroimage.2013.05.041. Epub 2013 May 16. PMID: 23684880; PMCID: PMC3724347.

Van Essen DC, Ugurbil K, Auerbach E, Barch D, Behrens TE, Bucholz R, Chang A, Chen L, Corbetta M, Curtiss SW, Della Penna S, Feinberg D, Glasser MF, Harel N, Heath AC, Larson-Prior L, Marcus D, Michalareas G, Moeller S, Oostenveld R, Petersen SE, Prior F, Schlaggar BL, Smith SM, Snyder AZ, Xu J, Yacoub E; WU-Minn HCP Consortium. The Human Connectome Project: a data acquisition perspective. Neuroimage. 2012 Oct 1;62(4):2222–31. doi: 10.1016/j.neuroimage.2012.02.018. Epub 2012 Feb 17. PMID: 22366334; PMCID: PMC3606888.

Vollenweider FX, Geyer MA. A systems model of altered consciousness: integrating natural and drug-induced psychoses. Brain Res Bull. 2001 Nov 15;56(5):495–507. doi: 10.1016/s0361-9230(01)00646-3. PMID: 11750795.

Wager TD, Atlas LY, Lindquist MA, Roy M, Woo CW, Kross E. An fMRI-based neurologic signature of physical pain. N Engl J Med. 2013 Apr 11;368(15):1388–97. doi: 10.1056/NEJMoa1204471. PMID: 23574118; PMCID: PMC3691100.

Wang Q, Wang Y, Kuo HC, Xie P, Kuang X, Hirokawa KE, Naeemi M, Yao S, Mallory M, Ouellette B, Lesnar P, Li Y, Ye M, Chen C, Xiong W, Ahmadinia L, El-Hifnawi L, Cetin A, Sorensen SA, Harris JA, Zeng H, Koch C. Regional and cell-type-specific afferent and efferent projections of the mouse claustrum. Cell Rep. 2023 Feb 11;42(2):112118. doi: 10.1016/j.celrep.2023.112118. Epub ahead of print. PMID: 36774552.

Wiech K, Kalisch R, Weiskopf N, Pleger B, Stephan KE, Dolan RJ. Anterolateral prefrontal cortex mediates the analgesic effect of expected and perceived control over pain. J Neurosci. 2006 Nov 1;26(44):11501–9. doi: 10.1523/JNEUROSCI.2568-06.2006. PMID: 17079679; PMCID: PMC2635565.

Wiech K, Seymour B, Kalisch R, Stephan KE, Koltzenburg M, Driver J, Dolan RJ. Modulation of pain processing in hyperalgesia by cognitive demand. Neuroimage. 2005 Aug 1;27(1):59–69. doi: 10.1016/j.neuroimage.2005.03.044. PMID: 15978845.

White MG, Cody PA, Bubser M, Wang HD, Deutch AY, Mathur BN. Cortical hierarchy governs rat claustrocortical circuit organization. J Comp Neurol. 2017 Apr 15;525(6):1347–1362. doi: 10.1002/cne.23970. Epub 2016 Feb 8. PMID: 26801010; PMCID: PMC4958609.

White MG, Mathur BN. Claustrum circuit components for top-down input processing and cortical broadcast. Brain Struct Funct. 2018 Dec;223(9):3945–3958. doi: 10.1007/s00429-018-1731-0. Epub 2018 Aug 14. PMID: 30109490; PMCID: PMC6252134.

White MG, Mathur BN. Frontal cortical control of posterior sensory and association cortices through the claustrum. Brain Struct Funct. 2018 Jul;223(6):2999–3006. doi: 10.1007/s00429-018-1661-x. Epub 2018 Apr 6. PMID: 29623428; PMCID: PMC5995986.

White MG, Mu C, Qadir H, Madden MB, Zeng H, Mathur BN. The Mouse Claustrum Is Required for Optimal Behavioral Performance Under High Cognitive Demand. Biol Psychiatry. 2020 Nov 1;88(9):719–726. doi: 10.1016/j.biopsych.2020.03.020. Epub 2020 Apr 10. PMID: 32456782; PMCID: PMC7554117.

White MG, Panicker M, Mu C, Carter AM, Roberts BM, Dharmasri PA, Mathur BN. Anterior Cingulate Cortex Input to the Claustrum Is Required for Top-Down Action Control. Cell Rep. 2018 Jan 2;22(1):84–95. doi: 10.1016/j.celrep.2017.12.023. PMID: 29298436; PMCID: PMC5779631.

Wilcox CE, Dekonenko CJ, Mayer AR, Bogenschutz MP, Turner JA. Cognitive control in alcohol use disorder: deficits and clinical relevance. Rev Neurosci. 2014;25(1):1–24. doi: 10.1515/revneuro-2013-0054. PMID: 24361772; PMCID: PMC4199648.

Worsley KJ, Marrett S, Neelin P, Vandal AC, Friston KJ, Evans AC. A unified statistical approach for determining significant signals in images of cerebral activation. Hum Brain Mapp. 1996;4(1):58–73. doi: 10.1002/(SICI)1097-0193(1996)4:1<58::AID-HBM4>3.0.CO;2-O. PMID: 20408186.

Xu QY, Zhang HL, Du H, Li YC, Ji FH, Li R, Xu GY. Identification of a Glutamatergic Claustrum-Anterior Cingulate Cortex Circuit for Visceral Pain Processing. J Neurosci. 2022 Oct 26;42(43):8154–8168. doi: 10.1523/JNEUROSCI.0779-22.2022. Epub 2022 Sep 13. PMID: 36100399; PMCID: PMC9637003.

Yao Z, Zhang Y, Lin L, Zhou Y, Xu C, Jiang T; Alzheimer’s Disease Neuroimaging Initiative. Abnormal cortical networks in mild cognitive impairment and Alzheimer’s disease. PLoS Comput Biol. 2010 Nov 18;6(11):e1001006. doi: 10.1371/journal.pcbi.1001006. PMID: 21124954; PMCID: PMC2987916.

Zang YF, He Y, Zhu CZ, Cao QJ, Sui MQ, Liang M, Tian LX, Jiang TZ, Wang YF. Altered baseline brain activity in children with ADHD revealed by resting-state functional MRI. Brain Dev. 2007 Mar;29(2):83–91. doi: 10.1016/j.braindev.2006.07.002. Epub 2006 Aug 17. Erratum in: Brain Dev. 2012 Apr;34(4):336. PMID: 16919409.

Zeidman P, Jafarian A, Corbin N, Seghier ML, Razi A, Price CJ, Friston KJ. A guide to group effective connectivity analysis, part 1: First level analysis with DCM for fMRI. Neuroimage. 2019 Oct 15;200:174–190. doi: 10.1016/j.neuroimage.2019.06.031. Epub 2019 Jun 19. PMID: 31226497; PMCID: PMC6711459.

Zeidman P, Jafarian A, Seghier ML, Litvak V, Cagnan H, Price CJ, Friston KJ. A guide to group effective connectivity analysis, part 2: Second level analysis with PEB. Neuroimage. 2019 Oct 15;200:12–25. doi: 10.1016/j.neuroimage.2019.06.032. Epub 2019 Jun 18. PMID: 31226492; PMCID: PMC6711451.

Zuo XN, Di Martino A, Kelly C, Shehzad ZE, Gee DG, Klein DF, Castellanos FX, Biswal BB, Milham MP. The oscillating brain: complex and reliable. Neuroimage. 2010 Jan 15;49(2):1432–45. doi: 10.1016/j.neuroimage.2009.09.037. Epub 2009 Sep 24. PMID: 19782143; PMCID: PMC2856476.

Zou QH, Zhu CZ, Yang Y, Zuo XN, Long XY, Cao QJ, Wang YF, Zang YF. An improved approach to detection of amplitude of low-frequency fluctuation (ALFF) for resting-state fMRI: fractional ALFF. J Neurosci Methods. 2008 Jul 15;172(1):137–41. doi: 10.1016/j.jneumeth.2008.04.012. Epub 2008 Apr 22. PMID: 18501969; PMCID: PMC3902859.

